# Mitochondria-containing extracellular vesicles from mouse *vs*. human brain endothelial cells for ischemic stroke therapy

**DOI:** 10.1101/2024.01.16.575903

**Authors:** Kandarp M. Dave, Venugopal R. Venna, Krithika S. Rao, Donna B. Stolz, Bodhi Brady, Victoria A. Quaicoe, Michael E. Maniskas, Ella E. Hildebrand, Dawson Green, Mingxi Chen, Jadranka Milosevic, Si-yang Zheng, Sruti S. Shiva, Louise D. McCullough, Devika S Manickam

## Abstract

Ischemic stroke-induced mitochondrial dysfunction in the blood-brain barrier-forming brain endothelial cells (**BECs**) results in long-term neurological dysfunction post-stroke. We previously data from a pilot study where *intravenous* administration of human BEC (**hBEC**)-derived mitochondria-containing extracellular vesicles (**EVs**) showed a potential efficacy signal in a mouse middle cerebral artery occlusion (**MCAo**) model of stroke. We *hypothesized* that EVs harvested from donor species homologous to the recipient species (*e.g.,* mouse) may improve therapeutic efficacy, and therefore, use of mouse BEC (**mBEC**)-derived EVs may improve post-stroke outcomes in MCAo mice.

We investigated potential differences in the mitochondria transfer of EVs derived from the same species as the recipient cell (mBEC-EVs and recipient mBECs or hBECs-EVs and recipient hBECs) *vs*. cross-species EVs and recipient cells (mBEC-EVs and recipient hBECs or *vice versa*). Our results showed that while both hBEC- and mBEC-EVs transferred EV mitochondria, mBEC-EVs outperformed hBEC-EVs in increasing ATP levels and improved recipient mBEC mitochondrial function via increasing oxygen consumption rates. mBEC-EVs significantly reduced brain infarct volume and neurological deficit scores compared to vehicle-injected MCAo mice. The superior therapeutic efficacy of mBEC-EVs in a mouse MCAo stroke support the continued use of mBEC-EVs to optimize the therapeutic potential of mitochondria-containing EVs in preclinical mouse models.

**Graphical Abstract:** 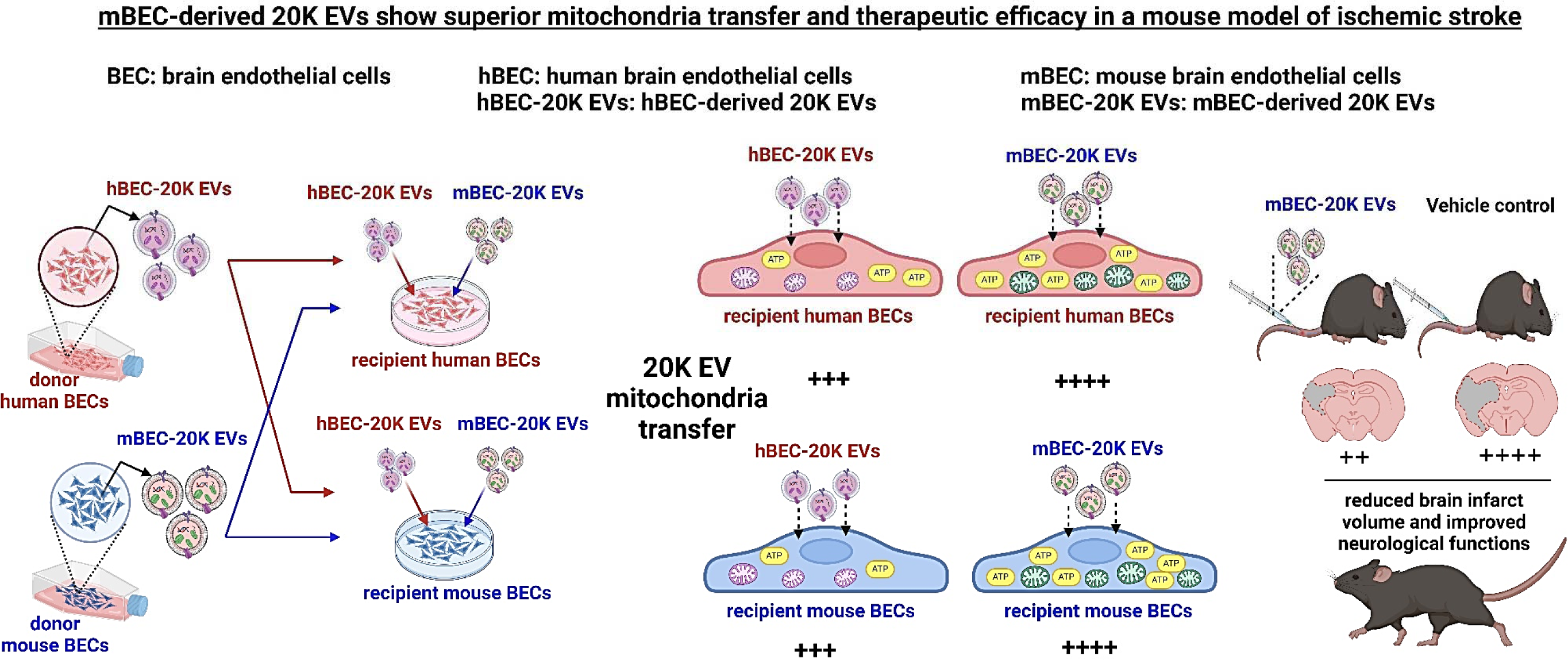

## 1. Introduction

Mitochondria are essential organelles that maintain cellular bioenergetics by producing adenosine triphosphate (**ATP**) and are involved in key metabolic pathways, enabling cell survival and proliferation [1–3]. Given their central role, mitochondrial damage or dysfunction produces excessive cytochrome c, reactive oxygen and nitrogen species, leading to programmed cell death. Mitochondrial dysfunction is one of the hallmarks of numerous pathological conditions [4], including ischemic stroke—the second leading cause of death worldwide [5, 6].

Ischemic stroke occurs when an obstruction in brain blood vessels reduces oxygen and nutrient supply to the affected brain regions, leading to brain damage, chronic disability, or even death [6, 7]. The blood-brain barrier (**BBB**)-forming brain endothelial cells (**BECs**) play a causal role in the pathophysiology of ischemic stroke. BECs contain about five-fold greater amounts of mitochondria than peripheral endothelial cells [8, 9]. The greater mitochondrial load in BECs is required to maintain cellular homeostasis and BBB integrity [3, 8, 10]. Oxygen-glucose deprivation (**OGD**) in BECs leads to excessive intracellular cation accumulation and glutamate excitotoxicity, impairing mitochondrial membrane potential. As a result, OGD-exposed BECs show reduced mitochondrial oxidative phosphorylation and ATP levels [10, 11], subsequently triggering cell death [12]. Therefore, maintaining BEC mitochondrial functionality is critical for cell survival. Physiologically, the damaged mitochondria are either rescued by fusion with intracellular healthy mitochondria or degraded by mitophagy—phagocytosis of damaged mitochondria by lysosomes [13]. When cells cannot recover damaged mitochondria under stress conditions, exogenous delivery of healthy mitochondria to the damaged cell could be a promising approach to mitigate mitochondrial dysfunction in ischemic stroke [14]. No clinically approved therapeutics deliver functional mitochondria for the treatment of mitochondrial disorders.

Numerous studies have shown that cell-derived extracellular vesicles (**EVs**) contain innate mitochondrial components such as mitochondrial DNA, proteins, and mitochondria [15–19] along with their intrinsic nucleic acids, non-mitochondrial proteins and lipid components. Moreover, the natural ability of EVs to protect their innate cargo from systemic degradation [19], cross the BBB [20], and deliver their innate cargo to the target cells [21] makes them a promising carrier for drug delivery to the brain [22, 23]. Exosomes (**EXOs**) and microvesicles (**MVs**) are subtypes of EVs that differ in their biogenesis, vesicle size, and innate components [18, 24, 25]. EXOs are derived from the inward budding of endosomal membranes and characteristic particle diameters of <200 nm. MV biogenesis involves outward blebbing and pinching of plasma membranes, releasing MVs into extracellular spaces with particle diameters >200 nm [26]. <u>In our study, EXOs and MVs were isolated at 20,000 ×g and 120,000 ×g, and are referred to as 20K EVs and 120K EVs, respectively, in line with the recommendations from the International Society of Extracellular Vesicles</u> [27, 28]. Published reports have shown that 120K EVs selectively contain mitochondrial proteins [15] and mitochondrial DNA [16], while 20K EVs contain mitochondrial proteins and functional mitochondria [18, 19]. Transfer of EV mitochondrial components from donor to recipient cells increased recipient cellular bioenergetics and cell survival in different experimental models of disease *in vitro* and *in vivo* [18, 29–31]. Importantly, delivery of mitochondria-containing EVs into recipient cells has resulted in a variety of therapeutic responses under various pathological conditions. For instance, Silva *et al.* demonstrated that 20K EV-mediated mitochondrial transfer into recipient pulmonary endothelial cells resulted in a three-fold increase in recipient cell ATP levels and a six-fold increase in mitochondrial respiration [32]. Our previous studies showed that human BEC cell line-derived 20K EVs contained mitochondria [33], that upon transfer into recipient human BECs increased cellular ATP levels, and mitochondrial respiration in normoxic and hypoxic culture conditions [33, 34]. Thus, delivery of mitochondria containing EVs is a promising treatment strategy to restore cellular bioenergetics in disorders associated with mitochondrial dysfunction.

The therapeutic efficacy of EVs depends on the source of EVs that may define its natural affinity to recipient cells and innate EV cargo. EV membranes may play a role in binding to the recipient cells and subsequent internalization into the target cells. EVs can be isolated from most eukaryotic cell types, and thus, selecting an appropriate EV donor cell type will likely influence the innate EV cargo and membrane characteristics that can ultimately determine the extent of therapeutic responses in the recipient cells. Harvesting EVs from the same type of donor cells as that of the target recipient cells may increase EV binding and internalization via “homotypic” interactions into the recipient cells. As BBB-lining BECs play a vital role in the pathophysiology of ischemic stroke, BEC-derived EVs may be an effective strategy to improve the functional integrity of recipient BECs and mitigate stroke-induced brain damage [35].

Moreover, the species of EV donor cells may contribute to the safety and efficacy of EV therapeutics in the preclinical and clinical phases of product development. Therapeutic EVs are often derived from human cell lines but are primarily tested in rodents and other animal disease models throughout preclinical development [36]. EV harvested from donor species different than the recipient species (“heterotypic” pair) may exert cross-species immunogenicity and toxicity to the recipient species and/or may compromise therapeutic efficacy [36]. Our previous pilot study administered human BEC (**hBEC**)-20K EVs via intravenous (**IV**) injection into a mouse middle cerebral artery occlusion (**MCAo**) model of ischemic stroke. In the absence of treatment-related mortalities, 20K EV-injected mice showed a trend of reduced brain infarct volume compared to vehicle-treated mice [33]. In this work, we *hypothesized* that *IV-*administration of mouse BEC (**mBEC**)-20K EV in a mouse MCAo model of stroke may exert superior therapeutic responses.

First, we isolated 120K EVs and 20K EVs from mBEC and hBEC cell lines and compared their particle characteristics, morphology and protein biomarkers. Second, we demonstrated the presence of mitochondrial components in hBEC and mBEC-20K EVs using transmission electron microscopy (**TEM**) and western blot analysis. Third, we evaluated homotypic and heterotypic EV donor cell-recipient cell pairs (1) to compare 20K EV mitochondrial transfer into recipient mBECs and hBECs under normoxic and OGD conditions using fluorescence microscopy and flow cytometry and (2) to determine the effect of EV exposure on recipient ischemic BEC ATP levels and mitochondrial respiration using a luminescence-based ATP and Seahorse mitochondrial function assays. Lastly, we determined the therapeutic effect of mBEC-20K EVs on the resulting brain infarct volume and neurological function in a mouse model of ischemic stroke. Our data demonstrated that mBEC-20K EVs outperformed hBEC-20K EVs in transferring the innate 20K EV mitochondrial load and concurrently mitochondrial function in recipient ischemic mBEC *in vitro*. Importantly, *IV*-injected mBEC-20K EVs showed a significant reduction in mouse brain infarct volume and significantly improved neurological functions in a mouse MCAo model of ischemic stroke—demonstrating cerebroprotection.

## 2. Materials and Methods

### 2.1. Materials

| **Material** | **Supplier** |
| --- | --- |
| Endothelial cell basal medium | Lonza (Walkersville, MD) |
| Pierce Micro BCA and BCA protein assay kits | Thermo Scientific (Rockford, IL). |
| Collagen Type I | Corning (Discovery Labware Inc., Bedford, MA) |
| Hydrocortisone, human basic fibroblast growth factor, ascorbic acid and rotenone | Sigma-Aldrich (Saint Louis, MO) |
| Penicillin-Streptomycin solution, Chemically Defined Lipid Concentrate | Invitrogen (Carlsbad, CA) |
| Polycarbonate centrifuge tubes | Beckman Coulter, Inc. (Brea, CA) |
| MitoTracker Deep Red FM | Thermo Scientific (Rockford, IL) |
| Mouse monoclonal antibodies against ATP5A (cat#ab14748), GAPDH (cat#ab8245) | Abcam (Cambridge, MA) |
| Mouse monoclonal antibody against CD9 (cat#ab92726) | Life Technologies Corporation (Eugene, OR) |
| Alexa Fluor 790-conjugated donkey anti-mouse IgG | Jackson ImmunoResearch Lab Inc. (West Grove, PA) |
| Cell Titer Glo 2.0 reagent | Promega (Madison, WI) |
| Radioimmunoprecipitation assay buffer | Alfa Aesar (Ward Hill, MA) |
| Nitrocellulose membrane | GE Healthcare Life Sciences (Germany) |
| Intercept blocking buffer | LI-COR Inc. (Lincoln, NE) |
| CellROX Reagent | Thermo Scientific (Rockford, IL) |

### 2.2. Cell culture

A mouse cerebral endothelial cell line (bEnd.3, cat# CRL2299) was purchased from ATCC (Manassas, VA) at passage number (P) 22 and used until P32 in all the experiments. ***<u>We refer to bEnd.3 cells as mBECs wherever applicable.</u>*** mBECs were cultured in complete growth medium containing glutamine-supplemented Dulbecco’s Modified Eagle’s Medium (DMEM/high glucose + Glutamax, Gibco, Carlsbad, CA) containing 4.0 mM of L-glutamine supplemented with 10% fetal bovine serum in tissue culture flasks. No antibiotics were added to the bEnd.3 cell culture medium. Pre-warmed complete growth medium was replenished every 48 hours until cells reached 80-90% confluency. For subculturing, the cells were washed with 1*x* phosphate buffer saline (**PBS**) and detached from the flask using TrypLE Express (Gibco, Denmark), followed by neutralization with complete growth medium before passaging or plating.

A human cerebral microvascular brain endothelial cell line (hCMEC/D3, cat#102114.3C) was received from Cedarlane Laboratories (Burlington, Ontario) at P25 and used between P25 and P35 in all experiments [33, 34, 37]. ***<u>We refer to hCMEC/D3 cells as hBECs wherever applicable.</u>*** hBECs were grown in tissue culture flasks and plates pre-coated using 0.15 mg/mL rat collagen I in a humidified 5% CO_2_ incubator at 37 ± 0.5°C (Isotemp, Thermo Fisher Scientific). The cells were cultured in complete growth medium composed of endothelial cell basal medium supplemented with 5% fetal bovine serum, penicillin (100 units/mL)-streptomycin (100 μg/mL) mixture (antibiotics was included here as recommended by the supplier), hydrocortisone (1.4 µM), ascorbic acid (5 µg/mL), Chemically Defined Lipid Concentrate (1% v/v), 10 mM HEPES (pH 7.4), and human basic fibroblast growth factor (1 ng/mL). The complete growth medium was replenished every 48 hours until the cells formed confluent monolayers. Cells were passaged as described in the previous paragraph. Cell suspensions stained with trypan blue (1:1 v/v ratio) were counted to calculate % live cells using a hemocytometer before passaging at 1:3 to 1:5 v/v ratios or plating in well plates at the indicated cell densities [38].

### 2.3. Isolation of 120K EVs and 20K EVs from mBEC and hBEC cell lines

We studied 120K EVs (exosomes, EXOs) and 20K EVs (microvesicles, MVs) separately and collectively refer to them as EVs wherever applicable. 20K EVs and 120K EVs were isolated from the conditioned medium of mBECs or hBECs using the differential ultracentrifugation method described earlier [33, 37, 38]. Briefly, mBECs and hBECs were cultured in complete growth medium in tissue culture flasks with a growth area of 175 cm^2^ (T175). Confluent BECs were washed with pre-warmed PBS pH 7.4 (0.0067M, PO_4_) without calcium and magnesium and incubated with serum-free medium for 48 h in a humidified incubator. Post-incubation, the EV-conditioned medium was collected in polypropylene tubes and centrifuged at 300 ×g for 11 min followed by 2000 ×g for 22 min to pellet down apoptotic bodies and cell debris using an Eppendorf 5810 R 15 amp centrifuge (Eppendorf, Germany). The supernatant was transferred to polycarbonate tubes and centrifuged at 20,000 ×g for 45 min at 4°C to pellet 20K EVs using a Sorvall MX 120+ micro-ultracentrifuge (Thermo Scientific, Santa Clara, CA). Next, the supernatant was filtered through 0.22 µm polyethersulfone filters into polycarbonate tubes and centrifuged at 120,000 ×g for 80 min to pellet 120K EVs. The supernatants were then discarded. 20K EVs and 120K EVs pellets were suspended in PBS pH 7.4 and stored at -20 °C until further use. The total EV protein content was measured using a Pierce micro BCA assay kit (Thermo Scientific, Rockford, IL) as described earlier [37, 38].

For isolation of mitochondria-depleted EVs, confluent mBEC and hBECs were treated with 0.25 μM rotenone (**RTN**, mitochondrial complex I inhibitor) in complete culture medium for four hours. Post-treatment, the treatment mixture was replaced with serum-free culture medium and incubated in a humidified incubator at 37 °C for 48 h. We isolated 120K EVs and 20K EVs from the conditioned medium of BECs using a differential ultracentrifugation method described above. EVs isolated post-RTN treatment were referred to as **RTN-120K EVs** and **RTN-20K EVs**, respectively.

For isolation of 20K and 120K blank medium controls, RTN at 100 nM was added to the respective serum-free, blank growth medium for hBECs or mBECs. RTN-containing medium was centrifuged at 300 ×g for 11 min at 4°C. The supernatant was transferred to a polypropylene tube and centrifuged at 2000 ×g for 22 min at 4°C. The supernatant was transferred to polycarbonate tubes and centrifuged at 20,000 ×g for 45 min at 4°C. The supernatant was transferred to another centrifuge tube, whereas the “apparent” pellet was suspended in PBS and is referred to as 20K-medium control. The supernatant was filtered through a 0.22 μm PES filter, transferred to polycarbonate tubes, and centrifuged at 120,000 ×g for 70 min at 4°C. The supernatant was discarded. The “apparent” pellet was suspended in PBS and is referred to as 120K-medium control.

### 2.4. Dynamic light scattering

Particle diameters and zeta potential of mBEC and hBEC-EVs were measured using dynamic light scattering. 20K EVs and 120K EVs were diluted to a final concentration of 0.1 mg EV protein/mL in either PBS pH 7.4 for particle diameters or in deionized water for zeta potential measurements. EV samples were analyzed using a Malvern Zetasizer Pro (Malvern Panalytical Inc., Worcestershire, UK). All samples were analyzed in triplicate measurements. Average particle diameter, dispersity index, and zeta potential values were reported as mean ± standard deviation (SD) of three independent experiments.

### 2.5. Nanoparticle tracking analysis

Stock samples of EVs (0.1 mg EV protein/mL) were diluted to 1:100 and 1:200 in PBS pH 7.4 and analyzed using a multiple-laser Zetaview f-NTA Nanoparticle Tracking Analyzer (Particle Metrix Inc., Mebane, NC). Prior to measurement, ZetaView was calibrated with 100 nm polystyrene beads and three 60 s videos were acquired at 520 nm laser wavelengths for particle diameter and concentration measurements. Additionally, a blank PBS sample was used prior to measuring the EV samples. In order to keep our Zetaview measurements uniform across extended periods, we used consistent materials, including fresh filtered PBS (Corning, Cat# 21-040-CV), 0.22 μm filter (Foxx Life Sciences, cat# 371-2215-OEM), and a syringe (Fisher Scientific, Cat# 14-955-457) to load samples. The methods we developed ensure that EV samples are measured without interference from external particles in PBS. Average EV concentrations were reported as mean ± SD of n = 3 measurements.

### 2.6. Transmission electron microscopy (TEM) of EVs

The morphology of negative-stained EVs and the presence of mitochondria in sectioned EVs were evaluated using the TEM described earlier [33].

#### Negative-stain images of EVs

EV suspensions (150 μL) were pelleted at 100,000 ×g in a Beckman Airfuge for 45 min; the supernatant was carefully removed, and the pellets were gently resuspended in 30 μL of PBS. Formvar coated, 100 mesh grids were floated onto 10 μL of this suspension and incubated for 5 min. The solution was wicked away with Whatman filter paper and stained with 0.45 µm filtered, 1% uranyl acetate. EV images were then acquired using a JEM-1400 Flash transmission electron microscope (JEOL, Peabody, 268 MA, USA) fitted with a bottom mount AMT digital camera (Danvers, MA, USA).

#### TEM of cross-sectioned EVs

EV suspensions were pelleted at 100,000 ×g in a Beckman airfuge for 20 min, and the pellets were fixed in 2.5% glutaraldehyde in PBS overnight. The supernatant was removed, and the cell pellets were washed 3× in PBS and post-fixed in 1% OsO4, 1% K_3_Fe(CN)_6_ for one hour. Following three additional PBS washes, the pellet was dehydrated through a graded series of 30–100% ethanol. After several changes of 100% resin over 24 h, the pellet was embedded in a final change of resin, cured at 37 °C overnight, followed by additional hardening at 65 °C for two more days. Ultrathin (70 nm) sections were collected on 200 mesh copper grids and stained with 2% uranyl acetate in 50% methanol for 10 min, followed by 1% lead citrate for seven min. Sections were imaged using a JEOL JEM 1400 Plus transmission electron microscope (Peabody, MA) at 80 kV fitted with a side mount AMT 2 k digital camera (Advanced Microscopy Techniques, Danvers, MA).

### 2.7. Western blot analysis

mBEC and hBEC-EVs were evaluated for characteristic protein biomarkers using western blotting. BEC lysates were used as positive controls. BEC lysate (10 µg) and 50 µg EV proteins were mixed with laemmli sodium dodecyl sulfate sample buffer and denatured at 95°C for 5 min. The samples were loaded onto a 4% stacking and 10% resolving polyacrylamide gel. Premixed molecular weight markers (Protein standards, ladder, 250 kDa - 10 kDa, cat#1610374, BioRad Laboratories Inc.) were used as a reference control to confirm molecular masses of the protein bands in the sample lanes. The gel was run at 120 V for 120 min in Tris-Glycine pH 8.8 running buffer using a PowerPac Basic setup gel electrophoresis assembly (BioRad Laboratories Inc.). The proteins were transferred onto a 0.45 µm nitrocellulose membrane at 75 V for 90 min using a transfer assembly (BioRad Laboratories Inc.). The membrane was washed with 0.1% Tween 20 containing Tris-buffered saline pH 7.4 (T-TBS) and blocked with Intercept blocking buffer solution (LI-COR Inc., Lincoln, NE) for one hour at room temperature. The membrane was incubated with mouse anti-GAPDH (1 μg/mL), mouse anti-ATP5A (1 μg/mL), rabbit anti-TOMM20 (1 μg/mL), rabbit anti-calnexin (1 μg/mL), rabbit anti-CD9 (0.2 μg/mL), rabbit anti-CD63 (1/1000 dilution) and rabbit anti-ARF-6 (1 μg/mL) primary antibodies in blocking buffer and T-TBS solution (1:1::v/v) overnight at 4°C. The membrane was washed with T-TBS and incubated with anti-mouse or anti-rabbit AF790 secondary antibodies (0.05 μg/mL) for one hour at room temperature. The membrane was washed with T-TBS and scanned using an Odyssey M imager (LI-COR Inc., Lincoln, NE) imager at 700 and 800 nm channels.

### 2.8. 20K EV mitochondria transfer into recipient BECs

#### 2.8.1. Isolation of mitochondria-labeled EVs

For labelling EV mitochondria, 120K EVs and 20K EVs were isolated from mBECs and hBECs prestained with Mitotracker deep red (**MitoT-red**). Confluent mBECs and hBECs in T175 flasks were stained with 100 nM MitoT-red in serum-free medium for 30 minutes in a humidified incubator. Post-incubation, the old medium was removed, and cells were washed with PBS pH 7.4. The cells were incubated in serum-free medium for 48 hours in a humidified incubator. Post-incubation, the conditioned medium was collected in polypropylene tubes. MitoT-red-20K EVs and MitoT-red-120K EVs were isolated using the differential ultracentrifugation method described in ***section 2.3***. The total protein content in MitoT-red-EVs was determined using a MicroBCA assay.

To isolate 20K and 120K-blank medium controls, Mitotracker deep red (at 100 nM) was added to the respective serum-free, blank growth medium for hBECs or mBECs. The dye-containing medium was centrifuged at 300 ×g for 11 min at 4°C. The supernatant was transferred to a polypropylene tube and centrifuged at 2000 ×g for 22 min at 4°C. The supernatant was transferred to polycarbonate tubes and centrifuged at 20,000 ×g for 45 min at 4°C. The supernatant was transferred to another centrifuge tube, whereas the “apparent” pellet was suspended in PBS and is referred to as 20K-medium control. The supernatant was filtered through a 0.22 μm PES filter, transferred to polycarbonate tubes, and centrifuged at 120,000 ×g for 70 min at 4°C. The supernatant was discarded. The “apparent” pellet was suspended in PBS and is referred to as 120K-medium control.

#### 2.8.2. Uptake of MitoT-red-EV into recipient mBECs and hBECs using fluorescence microscopy

EV mitochondria transfer into recipient BECs under normoxic conditions was studied using fluorescence microscopy. Recipient mBECs and hBECs were cultured in 24-well tissue culture plates at 100,000 cells/well in complete growth medium for 72 hours. The cells were treated with mBEC and hBEC-derived MitoT-red-EVs at 10, 30, and 60 μg EV protein/well in complete growth medium for 48 hours in a humidified incubator. Additionally, confluent recipient BECs were treated with 120K-medium control and 20K-medium control at equivalent volumes (used in the 10, 30, and 60 μg EV protein dose groups). Untreated cells incubated with complete growth medium were used as a control. Post-incubation, the treatment mixture was removed, and cells were washed with PBS. Cells were incubated with Hoechst stain (1 μg/mL) for 15 min. Cells treated with MitoT-red dye (100 nM for 30 min) were used as a positive control for MitoT-red signal detection. The cells were incubated in phenol red-free DMEM with 10% FBS medium prior to observation under an Olympus IX 73 epifluorescent inverted microscope (Olympus, Pittsburgh, PA) using Cyanine-5 (Cy5, excitation 651 nm and emission 670 nm) and DAPI channels at 20*x* magnification. Images were acquired and analyzed using CellSens Dimension software (Olympus, Pittsburgh, PA). The total sum of grayscale signal intensities in the Cy5 channel was used for signal quantification. The signal intensities of the untreated control group (background) were subtracted from signal intensities of the treated groups.

#### 2.8.3. Quantitative analysis of MitoT-red-EV mitochondria transfer into recipient mBECs and hBECs under normoxic and hypoxic conditions using flow cytometry

Recipient mBECs and hBECs were cultured in 24-well plates at 100,000 cells/well in complete growth medium. For normoxic conditions, confluent recipient BECs were treated with MitoT-red-120K EVs and MitoT-red-20K EV at 10, 30, and 60 μg EV protein/well in complete growth medium for 24 h in a humidified incubator. Additionally, confluent cells were treated with 120K-medium control and 20K-medium control at equivalent volumes (used in the 10, 30, and 60 μg EV protein dose groups). For OGD conditions, confluent cells were treated with MitoT-red-120K EVs and MitoT-red-20K EV at 10, 30, and 60 μg EV protein/well in oxygen-glucose deprived (**OGD**) medium in a hypoxia chamber (Billups Rothenberg, Del Mar, CA) pre-flushed with 5% carbon dioxide, 5% hydrogen and 90% nitrogen at 37±0.5°C for 24 h. The OGD medium [39] was composed of: 120 mM NaCl, 5.4 mM KCl, 1.8 mM CaCl_2_, 0.8 mM MgCl_2_, 25 mM Tris-HCl, pH 7.4. Unstained and untreated cells were used as a control. Post-treatment, the cells were washed with PBS, dissociated using TrypLE Express, diluted with PBS containing 1% FBS, and collected into centrifuge tubes. For each sample, an aliquot of a 100 μL cell suspension was analyzed through an Attune NxT Flow cytometer and 10,000 events were recorded in FSC *vs*. SSC plots. Mitotracker deep red fluorescence intensity was detected at 670/14 nm, and percentage signal intensities were analyzed in histogram plots generated using Attune 3.2.3 software. Mitotracker background signals were gated using the controls, including PBS and untreated cells.

We performed additional studies to determine Mitotracker red (**MitoT-red**)-EV mitochondria transfer into recipient hBECs at higher doses (>60 μg/well) using flow cytometry. We evaluated the MitoT-red-20K EV mitochondria transfer using the following parameters: 1) %MitoT+ve cells, which indicates the % population (*fraction*) of cells that internalize MitoT-red-20K EVs, and 2) mean fluorescence intensity (**MFI**), which represents the average fluorescence intensities of all cells within a sample, indicating the *magnitude* of MitoT-red transfer in MitoT+ve cells. Recipient hBECs were treated with hBEC- and mBEC-derived MitoT-red-20K EVs at 10, 30, 60, 90, and 120 μg EV protein/well in complete growth medium for 48 h in a humidified incubator. MitoT-red fluorescence intensity was detected at 670/14 nm, and the percentage of MitoT+ve cells and mean fluorescence intensities were analyzed in histogram plots generated using Attune 3.2.3 software.

### 2.9. Effect of EV treatment on relative ATP levels in the oxygen glucose-deprived recipient BECs

The effect of mBEC and hBEC-EVs on the relative cellular ATP levels in the recipient mBECs and hBECs under OGD conditions was measured using a Cell Titer Glo-based ATP assay. mBECs and hBECs in complete growth medium were cultured at 16,500 cells per well in a 96-well plate. The cells were incubated in a humidified incubator at 37°C for 72 h. Post-incubation, the cells were washed with PBS pH 7.4 and treated with OGD medium for four hours in an OGD chamber pre-flushed with 5% carbon dioxide, 5% hydrogen and 90% nitrogen at 37±0.5°C. Post-incubation, the OGD medium was removed, and cells were treated with mBEC- and hBEC-EVs diluted at different concentrations in the OGD medium for 24 h. Cells treated with OGD medium alone were used as OGD control, and cells treated with complete growth medium were used as normoxic control. After incubation, the treatment mixture was removed, and cells were washed and incubated with complete growth medium. An equal volume of Cell titer glo reagent was added. The plate was incubated for 15 min on a mechanical orbital shaker. A 60 µL aliquot of the mixture was transferred to a white opaque plate, and relative luminescence units at 1 s integration time were measured using a plate reader (Synergy HTX multimode plate reader, BioTek Inc., Winooski, VT). Relative ATP levels were calculated by normalizing the relative luminescence units (RLU) of treatment groups to the RLU of control, untreated cells.

We evaluated the effect of EVs isolated from donor cells with impaired mitochondrial complex I function on recipient BEC ATP levels. To inhibit mitochondrial function, EVs were isolated from the conditioned medium of mBECs and hBECs pretreated with 0.25 µM RTN, henceforth referred to as RTN-120K EVs and RTN-20K EVs. Confluent mBECs and hBECs in 96-well plates were treated with OGD medium for four hours in an OGD chamber pre-flushed with 5% carbon dioxide, 5% hydrogen and 90% nitrogen at 37±0.5°C. Post-incubation, the OGD medium was removed, and the cells were incubated with naïve 120K EVs, 20K EVs, RTN-120K EVs, and RTN-20K EVs-isolated from mBEC and hBECs at 10 μg EV protein/well under OGD conditions. Additionally, cells were treated with 120K- or 20K-medium controls at equivalent volume (used in the 10 μg EV protein dose groups) in OGD medium for 24 h. Normoxic BECs treated with complete culture medium and cells treated with OGD medium (OGD untreated cells) were used as controls. After incubation, the treatment mixture was removed, and the ATP assay was performed, as described in the paragraph above.

### 2.10. Measurement of mitochondrial function using Seahorse analysis

The mitochondrial function of recipient mBECs treated with EVs during OGD conditions were evaluated using the Seahorse analysis by measuring oxygen consumption (**OCR**) [34, 40]. mBECs seeded at 20,000 cells/well were cultured in a Seahorse XF96 plate for three days. Post-incubation, the cells were washed with PBS pH 7.4 and treated with OGD medium for four hours in a hypoxic chamber. The cells were incubated with mBEC and hBEC-derived 120K EVs and 20K EVs at different EV concentrations in OGD medium for 24 h. Post-incubation, the medium was replaced with pre-warmed DMEM and used for Seahorse analysis. After measurement of baseline OCR, 2.5 μmol/L oligomycin A and 0.7 μmol/L carbonyl cyanide-p-trifluoromethoxyphenyl-hydrazone were consecutively added to measure the proton leak and maximal OCR, respectively [40]. The total protein content of the cells in each well was measured using a Pierce BCA assay.

### 2.11. Cellular reactive oxygen species (ROS) measurements

Recipient hBECs and mBECs were cultured in black-walled and clear bottom tissue culture plates at 16,500 cells per well cell density in complete growth medium. At 90% confluency, the complete culture medium was removed, and cells were washed with 1x PBS and treated with OGD medium for four hours in an OGD chamber pre-flushed with 5% carbon dioxide, 5% hydrogen and 90% nitrogen at 37±0.5°C. Post-OGD exposure, the OGD medium was removed, and cells were treated with mBEC- and hBEC-EVs diluted at 10 μg EV protein/well in OGD medium for 24 h. Cells treated with OGD medium alone were used as OGD control, and cells treated with complete growth medium were used as normoxic control. After incubation, the treatment mixture was removed, and cells were washed with PBS. Cells were incubated with 5 μM CellROX deep red reagent diluted in complete growth medium for 30 min in a humidified incubator at 37°C. Post-incubation, the CellROX-containing medium was removed, and cells were washed thrice with 1x PBS. Cells in PBS were then measured to record fluorescence intensities at 640 nm excitation and 665 nm emission settings using a plate reader (Tecan Infinite M1000). Relative ROS levels were calculated by normalizing the fluorescence intensities of treated groups to those of control, untreated normoxic cells.

### 2.12. *In vivo* studies

#### 2.12.1. Mice

Young male C57BL/6 mice (8–12 weeks) were purchased from Jackson laboratories. Mice were acclimated in our animal facilities for one week before use. Mice were housed 4–5 per cage, with a 12-h light/dark schedule in a temperature and humidity-controlled vivarium, with *ad-libitum* access to a pelleted rodent diet (irradiated LabDiet 5053, PicoLab rodent diet 20) and filtered water. All experiments were approved by the Institutional Animal Care and Use Committee at the University of Texas Health Science Center, Houston. This study was performed in accordance with the guidelines provided by the National Institute of Health and followed RIGOR guidelines. Only male animals were used in this study to allow comparisons to earlier studies [33] and to exclude potential sex and estrogen-related effects. Animals were randomly assigned to treatment groups.

#### 2.12.2. Mouse middle cerebral artery occlusion stroke model

Focal transient cerebral ischemia was induced in mice by right middle cerebral artery occlusion (**MCAo**) followed by reperfusion as described previously [41, 42]. Briefly, young (3-month) male C57BL/6 mice were initially placed and anaesthetized in a chamber with 4% isoflurane, and adequate sedation was confirmed by tail pinch for all surgical procedures. Surgeries were performed under 1–2% continuous isoflurane. A 90-minute MCAo was achieved by blocking blood flow to MCA using a silicone-coated filament via the external carotid artery. At the end of ischemia (90 min MCAo), mice were briefly re-anesthetized, and reperfusion was initiated by filament withdrawal. Body temperature was maintained by placing the mouse on a heating pad at ∼37 ◦C during surgical procedures. Two hours after stroke onset, mice were randomly assigned to receive 100 μg of mBEC-20K EVs in 200 μL PBS (n=7) or vehicle (PBS, n=8) treatment by intravenous tail vein injection. Sample size was determined using power analysis based on results from the initial cohort [33]. Mice were euthanized 24 h after the stroke, and brains were analyzed for infarct size using 2,3,5-triphenyl tetrazolium chloride (**TTC**)-stained sections. Infarct analysis was performed by an investigator blinded to the treatment groups.

##### Details on the numbers of animals per group

We initially started with 18 mice for stroke surgeries, but one mouse did not show a reduction of blood flow to less than 70% from baseline laser Doppler flowmetry (**LDF**) values during middle cerebral artery occlusion (**MCAo**), and that mouse was excluded even before assigning to any treatment. We use LDF reductions as inclusion/exclusion criterion for further use of the MCAo mice. A less than 70% LDF reduction from baseline is used to determine if the mouse has “truly experienced a stroke”. The remaining mice were randomly assigned: 9 to PBS and 8 to EV treatment. A total of two mice died following MCAo surgeries, with one each from PBS and EV-treated groups.

#### 2.12.3. Neurological Deficit Scoring (NDS)

The behavioral functions of MCAo mice was examined by a blinded investigator using the Neurological Deficit Score (NDS) at 24 h after stroke. We used our standard graded scoring system to get NDS as follows: 0 = no deficit; 1 = forelimb weakness and torso turning to the ipsilateral side when held by the tail; 2 = circling to affected side; 3 = unable to bear weight on affected side; and 4 = no spontaneous locomotor activity or barrel rolling, as previously described [43–47].

### 2.13. Statistical analysis

Statistically significant differences between the means of controls and treatment groups or within treatment groups were determined using one-way or two-way analysis of variance (ANOVA) with post hoc corrections using multiple comparison tests at α=0.05 using GraphPad Prism 10 (GraphPad Software). Mann–Whitney U test was used for ordinal data (i.e., NDS) analysis. The notations for the different significance levels are indicated as follows: *p < 0.05, **p < 0.01, ***p < 0.001, ****p < 0.0001.

## 3. Results

### 3.1. Physicochemical characterization of mouse and human BEC-derived EVs

We used b.End3 and hCMEC/D3 cells as models of mouse (**mBEC**) and human BECs (**hBEC**), respectively. We compared mBEC-*vs.* hBEC-EVs in terms of their physicochemical characteristics, morphology, and EV biomarkers. Average particle diameters of mBEC 120K EVs and 20K EVs were 167 nm and 442 nm, with characteristic negative zeta potentials of about -25.1 mV and -27.5 mV, respectively (**Fig. 1a**). hBEC-120K EVs and 20K EVs showed average particle diameters of 137 nm and 214 nm with average zeta potentials of -13.4 mV and -24.3 mV, respectively (**Fig. 1a**). For both cell lines, 20K EVs showed significantly greater particle sizes than 120K EVs, and there was no significant difference between the average particle diameters of mBEC and hBEC-120K EVs. However, mBEC-20K EVs showed significantly greater particle diameters compared to hBEC-20K EVs. Notably, mBECs secreted EVs at lower concentrations of about 5.1×10^7^ particles/mL 20K EVs and 2.8×10^8^ particles/mL 20K EVs (**Fig. 1a**). In contrast, hBEC secreted about 2.1×10^9^ particles/mL 120K EVs and 1.7×10^9^ particles/mL 20K EVs (**Fig. 1a**).

**Figure 1:**
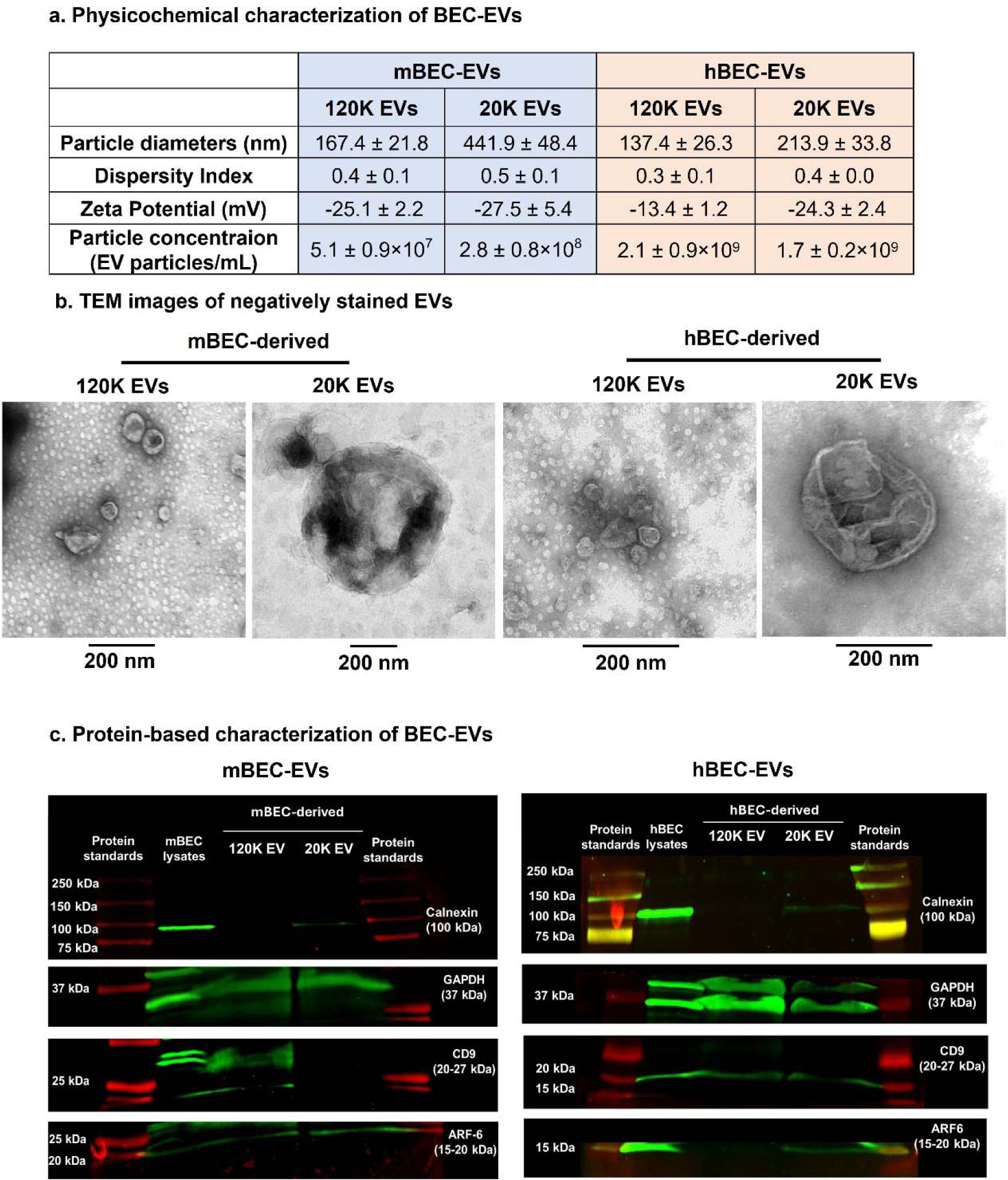
Physicochemical characterization of mBEC and hBEC-EVs. **a)** *Dynamic light scattering (DLS) and nanoparticle tracking analysis (NTA) of mBEC and hBEC-120K EVs and 20K EVs.* Freshly isolated 120K EVs and 20K EVs were diluted to 0.1 mg EV protein/mL in PBS pH 7.4 for particle diameters and NTA. The samples were diluted in deionized water for zeta potential measurements. Dynamic light scattering analysis of EVs was conducted on Malvern Zetasizer Pro-Red (Malvern Panalytical), and EV particle concentrations were measured on a multiple-laser ZetaView f-NTA Nanoparticle Tracking Analyzer (Particle Metrix Inc., Mebane, NC). DLS data are presented as mean ± SD of n = 3 independent experiments, and NTA data are presented as mean ± SD of n=3 measurements. **(b)** *Morphology of EVs.* Representative negative stain transmission electron microscopy (TEM) images of mBEC and hBEC-EVs. Scale bar: 200 nm. **(c)** *Detection of membrane and luminal biomarkers of mBEC and hBEC-derived EVs using western blot analysis.* Western blot shows the relative expression of calnexin (100 kDa), glyceraldehyde 3-phosphate dehydrogenase (GAPDH, 37 kDa), CD9 (20-27 kDa), and ARF6 (15-20 kDa) in 120K EVs and 20K EVs. Cell lysates were used as a control, and protein standards were used as a reference for the molecular weight of protein bands. Cell lysates (15 μg) and EV lysates (50 μg) mixed with laemmli sample buffer were electrophoresed on a sodium dodecyl sulfate-polyacrylamide gel. The proteins were transferred onto a nitrocellulose membrane, and non-specific proteins were blocked using a blocking buffer. The blot was incubated with primary antibodies overnight at 4 °C and subsequently incubated with secondary antibodies at room temperature. The blots were imaged on the 700 and 800 nm channels using an Odyssey M imager (LI-COR Inc. Lincoln, NE) and processed using ImageStudio 5.2 software.

Representative TEM images of negatively-stained EVs derived from both cell lines showed that 120K EVs were <200 nm and 20K EVs were >200 nm with heterogenous shapes (**Fig. 1b**). The observed morphologies are consistent with TEM images of EVs in the published literature [48, 49]. The larger field of **Fig. 1b** of mBEC- and hBEC-20K EVs, are now shown in **SL Fig. 1**. Additionally, we have acquired multiple TEM images of negatively-stained EVs (**SL Fig. 2**) to present the diverse morphology of 20K EVs. As anticipated, mBEC and hBEC 20K EVs exhibited various sizes and shapes. Notably, all hBEC 20K EVs were approximately 200 nm in diameter (**SL Fig. 2**). In contrast, mBEC 20K EVs were predominantly larger than 200 nm (**SL Fig. 2**). The representative TEM images of mBEC-20K EVs in the revised **Fig. 1b** show significantly larger particle diameters compared to hBEC-20K EVs, aligning with DLS data in **Fig. 1a**.

We compared membrane and luminal biomarker proteins in mBEC *vs*. hBEC-EVs using western blot analysis. According to the Minimal Information for Studies of Extracellular Vesicles (MISEV) 2018 guidelines, protein content-based characterization of EVs should demonstrate the purity and characteristic proteins associated with EV subtypes. We performed western blot analysis of mBEC and hBEC-120K EVs and 20K EVs to evaluate the relative expression of calnexin (to demonstrate that EVs are free of endoplasmic reticulum contaminants), tetraspanin CD9 and CD63 (transmembrane proteins associated with the plasma membrane or endosome), and glyceraldehyde 3-phosphate dehydrogenase (GAPDH as a model cytosolic protein recovered in EVs). mBEC and hBEC cell lysates showed bands of all the above proteins at their characteristic molecular masses (**Fig. 1c**), suggesting the suitability of western blotting for detecting these biomarker proteins and their relative abundance in the donor BECs. First, calnexin (endoplasmic reticulum marker, 100 kDa) was used to determine whether isolated EV samples contained cellular contaminants or non-EV co-isolated structures. mBEC- and hBEC-120K EVs showed no calnexin expression, suggesting that isolated 120K EVs were free from endoplasmic reticulum contaminants. In contrast, faint expression of calnexin in 20K EVs (**Fig. 1c**) indicated minimal endoplasmic reticulum impurities in the isolated 20K EVs. Second, mouse and human BEC-120K EVs and 20K EVs expressed GAPDH, suggesting the incorporation of soluble cytosolic proteins in EVs during their biogenesis. Third, tetraspanin CD9 (20-27 kDa) was selectively present in mBEC-120K EVs, whereas mBEC-20K EVs did not show the CD9 band (**Fig. 1c**). However, CD9 was detected in both hBEC-120K EVs and 20K EVs. Lastly, mBEC and hBEC-20K EVs showed adenosine diphosphate-ribosomal factor (ARF6) expression at its characteristic molecular weight of 15-20 kDa. In contrast, mBEC-120K EVs showed a faint band, and hBEC-120K EVs did not show any ARF6 band (**Fig. 1c**). Both mBEC- and hBEC-20K EVs expressed ARF6 protein.

### 3.2. Mouse and human BEC-20K EVs contained mitochondria and mitochondrial proteins

We acquired TEM images of cross-sectioned 120K EVs and 20K EVs isolated from mBEC and hBECs (**Fig. 2a**). The yellow arrowheads point to 120K EVs membranes, the blue arrowhead indicates 20K EVs membranes and the maroon arrowheads point to 20K EVs mitochondria. Mitochondria were detected as membrane-bound electron-dense structures in the 20K EVs lumen. Representative TEM images showed that mBEC- and hBEC-20K EVs (blue arrows, **Fig. 2a**) contained a couple of mitochondria (maroon arrows) with varying sizes and morphologies in the 20K EV lumen. In contrast, 120K EVs lumen lacked electron-dense structures, suggesting the absence of mitochondria in mBEC and hBEC-120K EVs. It is important to note that the morphology of 20K EVs mitochondria in our TEM images was comparable with extracellular free mitochondria or mitochondria-containing EVs presented in previously published reports [17, 19, 50]. So, we conclude that irrespective of the donor cell species, both mBEC and hBEC-20K EVs contained mitochondria, and 120K EVs lacked mitochondria.

**Figure 2:**
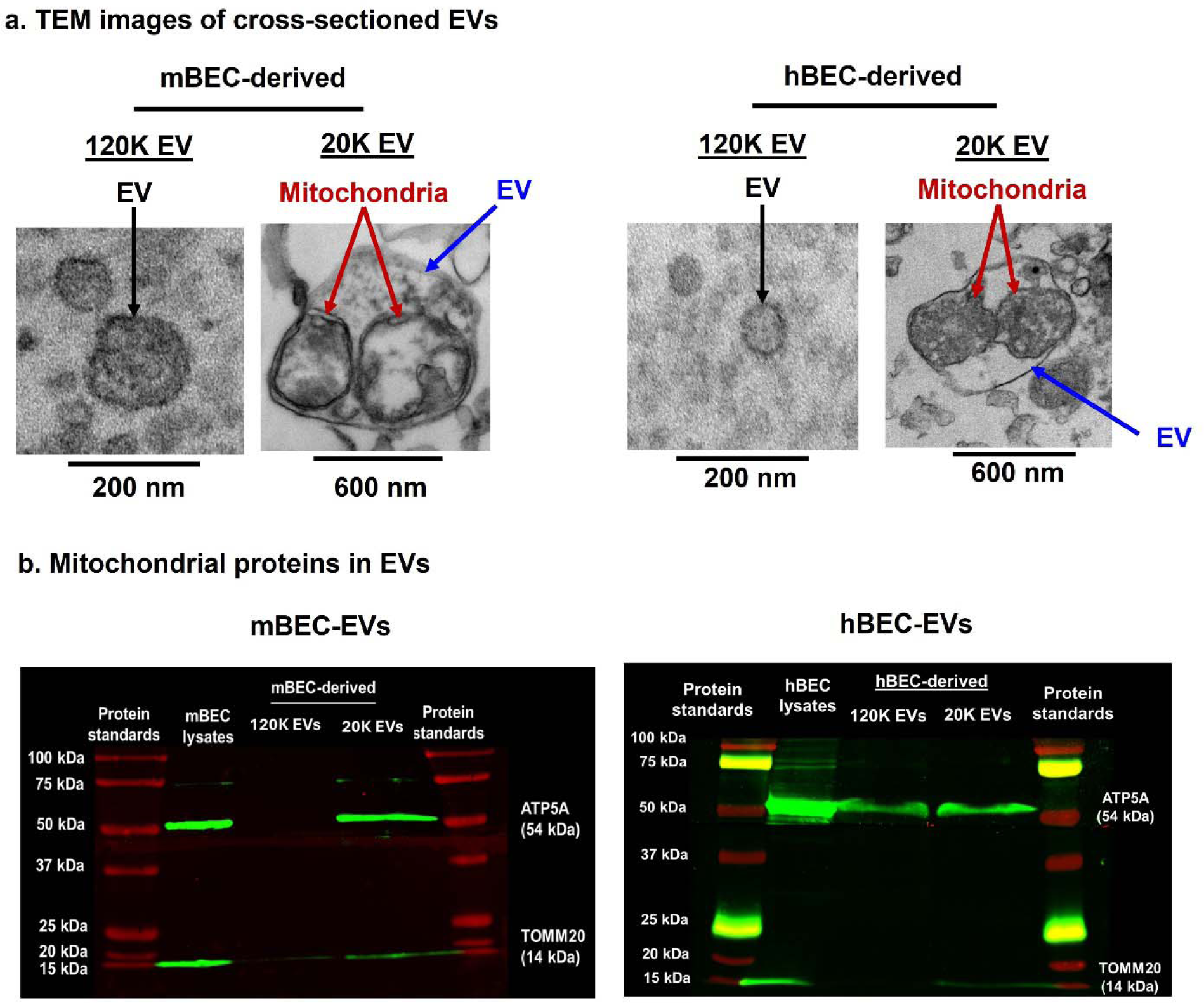
Selective presence of mitochondria in 20K EVs. (**a**) *Representative TEM images of sectioned mBEC and hBEC-120K EVs and -20K EVs.* mBEC- and hBEC-20K EVs (blue arrow) contained a couple of mitochondria (electron-dense structures, maroon arrows) in the 20K EV lumen. 120K EV (yellow arrows)-derived from both cell lines lacked electron-dense structures in the lumen. (**b**) *Detection of mitochondrial proteins in mBEC- and hBEC-EVs using western blot analysis.* Western blots show the relative expression of ATP5A (54 kDa) and TOMM20 (14 kDa) in 120K EVs and 20K EVs. Cell lysates were used as a control, and protein standards were used as a reference for the molecular mass of protein bands. The blots were imaged on the 700 and 800 nm channels using an Odyssey M imager (LI-COR Inc. Lincoln, NE) and processed using ImageStudio 5.2 software.

We further evaluated if structural and functional mitochondrial proteins are expressed in 120K EVs and 20K EVs using western blot analysis. Mouse and human BEC-120K EVs and 20K EVs were analyzed for the expression of ATP5A (a subunit of electron transport chain ATP synthase, 60 kDa) and TOMM20 (translocase of the outer mitochondrial membrane complex subunit 20, 14 kDa). While mBEC-20K EVs showed the presence of ATP5A and TOMM20 at their characteristic molecular masses, 120K EVs did not show any mitochondrial protein bands (**Fig. 2b**). The selective presence of mitochondrial proteins in 20K EVs but not in 120K EVs aligned with the TEM images of sectioned mBEC-20K EVs showing mitochondria in 20K EVs (**Fig. 2a**). However, ATP5A was detected in both 120K EVs and 20K EVs derived from hBECs. Importantly, TOMM20 was selectively present in hBEC-20K EVs (**Fig. 2b**), whereas 20K EVs showed no bands at the characteristic molecular mass of 14 kDa. It is likely that while both hBEC-120K EVs and 20K EVs contained functional mitochondrial proteins (ATP5A), and mitochondrial structural components (TOMM20) were selectively present in 20K EVs, aligning with the TEM images of sectioned hBEC-20K EVs showing mitochondria (**Fig. 2a**).

### 3.3. Mouse BEC-20K EVs transferred mitochondria into recipient mouse and human BECs

We sought to determine whether mBEC-20K EVs can transfer their innate mitochondria into recipient hBECs and *vice versa* (i.e., if hBEC-20K EVs can transfer their innate mitochondria into recipient mBECs) using fluorescent microscopy. Mitochondria-labelled EVs were isolated from donor mBECs and hBECs-prestained using Mitotracker Red (**MitoT-red-EVs**). Subsequently, recipient mBECs and hBECs were incubated with MitoT-red-EVs at different doses, and we detected MitoT-red signals 48 h-post exposure using a fluorescence microscope. MitoT-red is a mitochondrial membrane potential dye that stains polarized (functional) mitochondria in donor cells. Untreated mBECs and hBECs showed no signals in the Cy5 channel. Cells stained with MitoT-red showed intracellular purple fluorescence signals (**Fig. 3a-b**), suggesting minimal background and specificity of MitoT-red signal detection under the Cy5 channel. mBEC- and hBEC-derived MitoT-red-120K EVs showed faint MitoT-red signals at 10, 30, and 60 μg doses in recipient mBECs and hBECs (**Fig. 3a-b**). While MitoT-red-120K EVs at 10 and 30 μg doses exhibited detectable signals, the intensity is less pronounced compared to the 60 μg dose. The data suggests that 120K EVs likely contain minimal mitochondrial load and show signs of low transfer at higher doses. The quantitative analysis of MitoT-red signals (**Fig. 3c-d**) showed that mBEC-120K EVs showed a dose-dependent gradual increase in MitoT-red signals in recipient mBECs, whereas increasing the dose of hBEC-120K EVs did not increase MitoT-red signals. Notably, mBEC-120K EVs showed relatively higher MitoT-red signals (**Fig. 3c-d**), suggesting mBEC-120K EVs likely contain a greater mitochondrial load than hBEC-120K EVs.

**Figure 3a-b:**
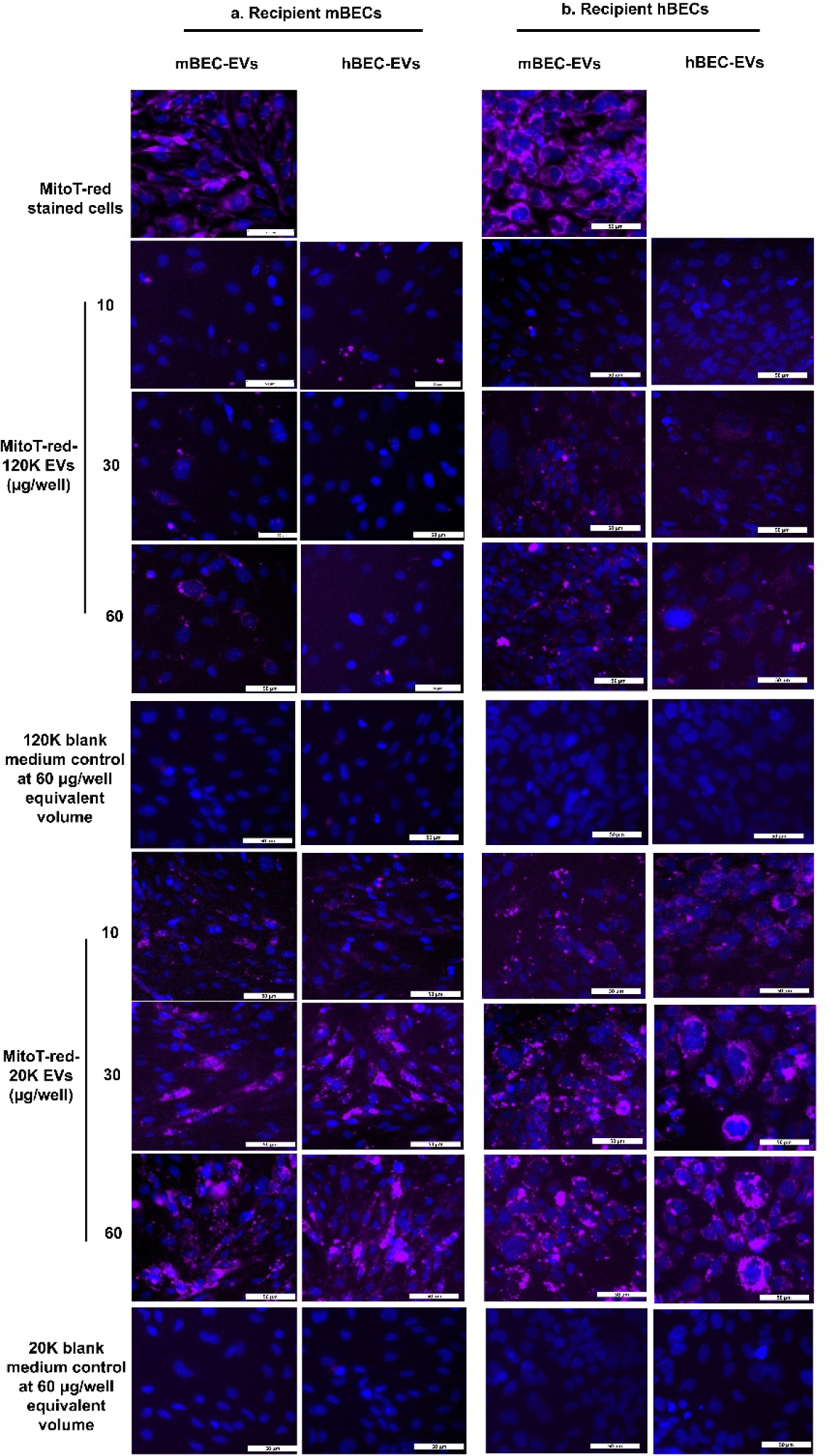
20K EV-mediated mitochondria transfer into the recipient BECs at varying doses. Recipient mBECs (**a**) and hBECs (**b**) cultured in 48-well plates were incubated with indicated amounts of mBEC and hBEC-derived MitoT-red-120K EVs, 20K EVs, and blank medium controls diluted in complete growth medium for 48 h. Cells treated with complete growth medium were used as control, untreated cells. Cells treated with 100 nM MitoT-red for 30 min were used as a positive control for MitoT-red signal detection under a fluorescence microscope. Post-incubation, the cells were washed and incubated with the phenol-red medium. MitoT-red signals (purple puncta) and nucleus (blue) were observed under an Olympus IX 73 epifluorescent inverted microscope using Cy5 (pseudo purple colored for mitochondria) and DAPI (pseudo blue colored for nucleus) channels at 20× magnification—scale bar 50 μm.

**Figure 3c-d:**
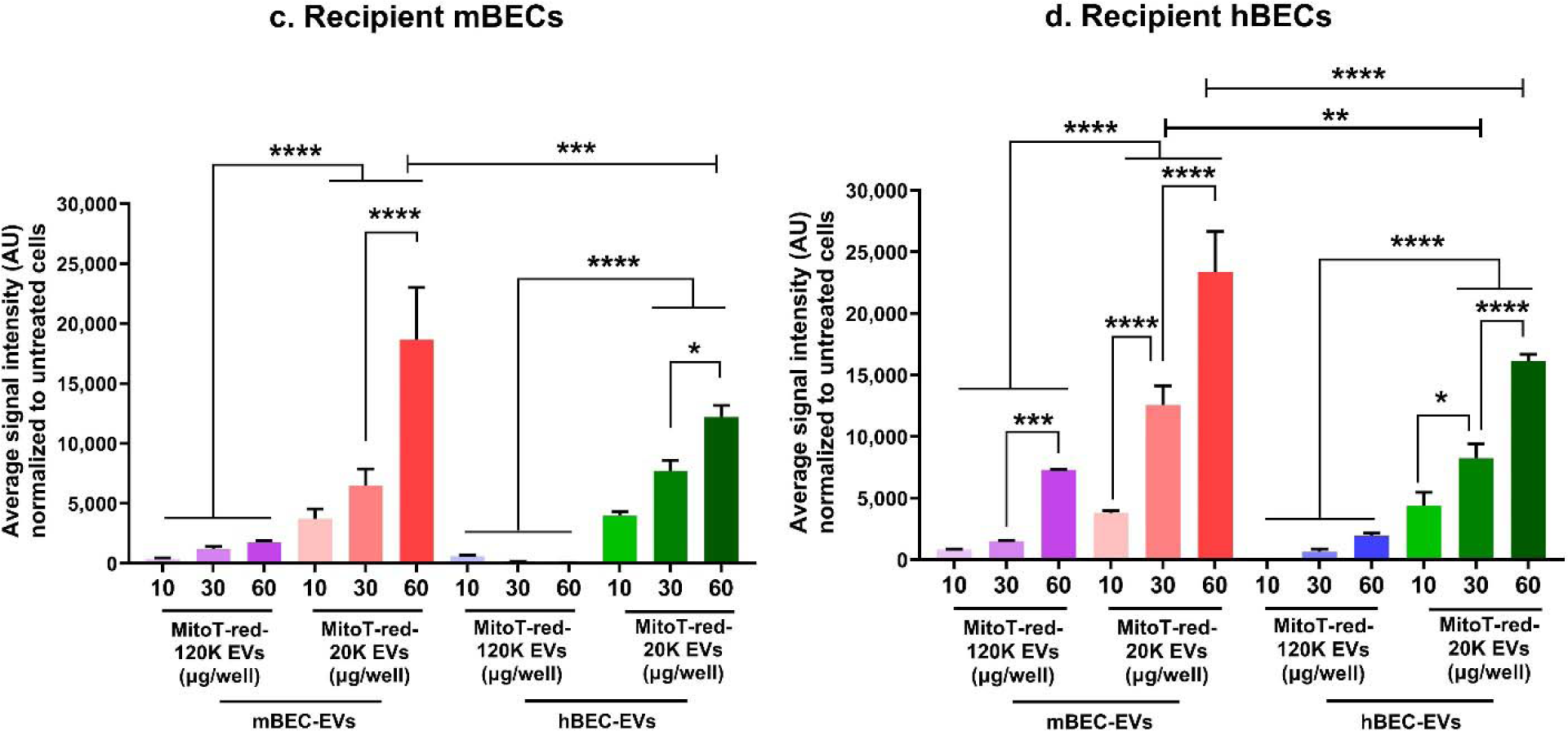
Quantification of 20K EV mitochondria transfer in recipient BECs. mBECs (**c**) and hBECs (**d**) were treated with mBEC and hBEC-derived MitoT-red-120K EVs and 20K EVs at indicated doses in complete growth medium for 48 h. Post-incubation, at least three images were acquired from each treatment group using an Olympus IX 73 epifluorescent inverted microscope. The total sum of grayscale signal intensities in the Cy5 channel was estimated using Olympus CellSens software. The signal intensities of the untreated control group (background) were subtracted from the signal intensities of the treatment groups. For the quantification of MitoT-positive cells, we used a total of 3 images per group (n=3). Each image contained an average of approximately 300-400 cells, resulting in an analysis of around 900-1200 cells per group. Data are presented as mean ± SD (n=3 images per treatment group). The statistically significant difference between treatment groups was analyzed using a one-way analysis of variance followed by post hoc Tukey’s multiple comparison test using GraphPad Prism 10. *p < 0.05, **p < 0.01, ***p < 0.001, ****p < 0.0001.

Recipient mBECs and hBECs treated with mBEC- and hBEC-derived MitoT-red-20K EVs at 10 μg showed strong intracellular MitoT-red signals (**Fig. 3a-b**). Greater doses (30 and 60 μg EV protein/well) of MitoT-red-20K EVs showed significantly higher intracellular MitoT-red signals in both recipient mBECs and hBECs (**Fig. 3a-b**). Importantly, quantitative analysis of MitoT-red signals showed that recipient BECs treated with 20K EVs at a dose as low as 10 μg led to a considerably greater extent of mitochondria transfer than the maximum dose of 120K EVs (**Fig. 3c-d**), suggesting a significantly greater mitochondrial load in 20K EVs than 120K EVs. Moreover, increasing 20K EVs doses showed a dose-dependent and statistically significant (p<0.0001) increase in recipient BEC signals (**Fig. 3c-d)**. Notably, there was no statistically significant difference between mBEC and hBEC-20K EVs -mediated mitochondria transfer at the low and medium 20K EVs doses in both recipient BECs. However, at higher doses, mBEC-20K EVs showed a significantly greater mitochondria transfer than hBEC-20K EVs in both recipient BECs (**Fig. 3c-d**), likely because of a greater mitochondrial load in mBEC-20K EVs than hBEC-20K EVs.

Next, we used flow cytometry to analyze 20K EVs mitochondria transfer into recipient BECs and determine whether ischemic conditions can affect 20K EV mitochondria transfer into oxygen-glucose-deprived (**OGD**) recipient BECs. First, recipient mBECs and hBECs were treated under normoxic conditions with 10, 30, and 60 μg of MitoT-red-EVs derived from mBECs or hBECs for 24 h (**Fig. 4a-d**). Recipient mBECs treated with different doses of mBEC-derived MitoT-red-120K EVs and 20K EVs showed a dose-dependent, statistically significant (p<0.0001) increase in MitoT-red positive cells (**Fig. 4a**). Consistent with microscopy studies (**Fig. 3c-d**), 20K EV-treated BECs showed significantly (p<0.0001) higher MitoT-red positive cells than 120K EVs-treated cells (**Fig. 4a**). Similarly, recipient mBECs treated with hBEC-derived MitoT-red-EVs (**Fig. 4b**) and recipient hBECs treated with mBEC- (**Fig. 4c**) as well as hBEC-derived MitoT-red-EVs (**Fig. 4d**) showed (1) a dose-dependent increase in MitoT-red positive cells, and (2) significantly greater extent of mitochondria transfer from 20K EVs than 120K EVs. Importantly, recipient mBECs treated with mBEC-derived MitoT-red-EVs under OGD conditions showed a dose-dependent, significant (p<0.0001) increase in MitoT-red positive cells (**Fig. 4a**), suggesting that 20K EVs mitochondria were internalized by ischemic BECs. However, recipient mBECs treated with MitoT-red-EVs showed significantly (p<0.0001) reduced MitoT-red positive cells under OGD conditions compared to normoxic cells treated at the same dose (**Fig. 4a**), suggesting that OGD reduces BEC efficiency to internalize EV mitochondria than normoxic conditions. Notably, at the highest treated dose, the homotypic 20K EV donor-recipient BEC pair (**Fig. 4a** and **c**) did not show any significant difference in percentage MitoT +ve cells between normoxic and OGD conditions, likely due to saturation of 20K EV mitochondria transfer into recipient normoxic BECs at the highest tested dose. The OGD-mediated reduction in EV mitochondria uptake in BECs compared to normoxic conditions was consistent in mouse and human recipient BECs, irrespective of the EV source species (**Fig. 4b-d**).

**Figure 4:**
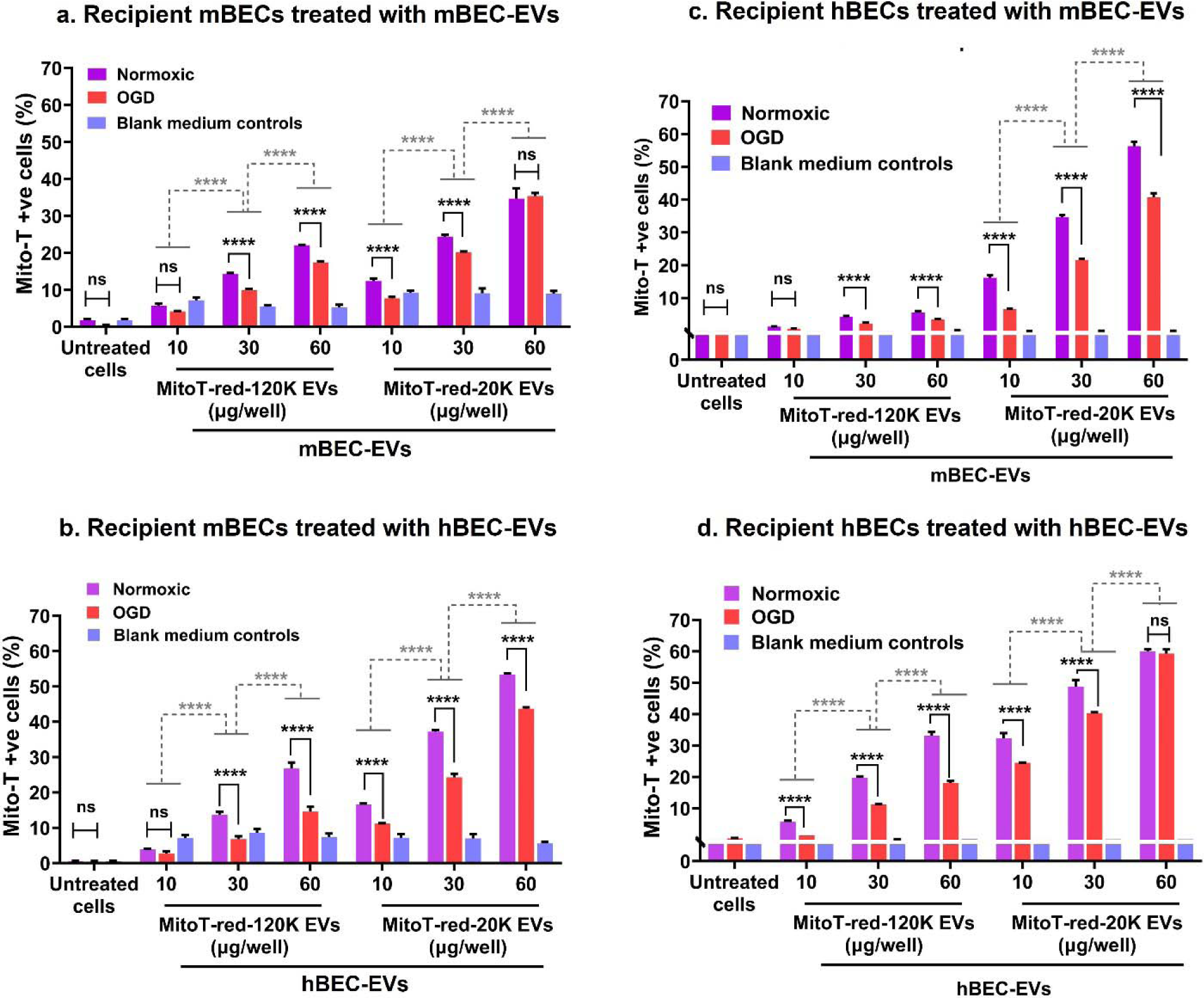
Transfer of EV mitochondria into recipient BECs at varying doses. Confluent mBECs and hBECs were incubated with the indicated amounts of mBEC and hBEC-derived MitoT-red-120K EVs, 20K EVs, and blank medium controls under normoxic and OGD conditions for 24 h. Post-incubation, the cells were washed, collected, and run through an Attune NxT flow cytometer. The histograms of mBECs and hBECs treated at indicated doses of MitoT-red-EVs were collected using a 674/10-nm side scatter filter in the Attune flow cytometer. Untreated and unstained BECs were used as controls to gate the background signals in histograms. MitoT-red-stained BECs were used as a positive control to gate the histograms for MitoT-red-positive counts. Subsequently, this gate was applied to quantify the percentage of MitoT-red BECs treated with MitoT-red-EVs. The percentage of maximum signal intensities of mBECs treated with mBEC-EVs (**a**), mBECs treated with hBEC-EVs (**b**), hBECs treated with mBEC-EVs (**c**), and hBECs treated with hBEC-EVs (**d**), were compared under normoxic and hypoxic conditions. Data represent mean ± SD of n = 3. The statistically significance between treatment groups was analyzed using one-way ANOVA followed by post hoc Tukey’s multiple comparison test using GraphPad Prism 10. *p < 0.05, **p < 0.01, ***p < 0.001, ****p < 0.0001.

Recipient mBECs treated with either 120K or 20K-blank medium controls showed lower than 10% of MitoT-positive cells at lower dose volumes (**Fig. 4a**). Increasing the dose from 10 to 60 μL for 120K- or 20K-medium controls did not result in any increase in MitoT-positive cells (**Fig. 4a**). Similarly, recipient mBECs treated with low, medium, and high doses of 120K and 20K-blank medium controls showed less than 1% MitoT-positive cells and did not show volume increment-mediated increase in MitoT-positive cells (**Fig. 4b**). Consistent with these findings in recipient mBECs, recipient hBECs treated with blank medium controls at low, medium, and high dosing volumes also showed less than 1% MitoT-positive cells and also did not show volume increment-mediated increase in MitoT-red-positive cells (**Fig. 4c** and **4d**).

Additionally, we performed microscopy studies to determine Mitotracker signals in recipient hBEC and mBECs treated with 20K or 120K-blank medium controls for 48 h (**Fig. 3a** and **b**). Consistent with the flow cytometry data, Our microscopy studies confirmed that neither the 20K nor blank medium controls showed any significant MitoT-red signals at low, medium, and high dosing volumes, indicating that the observed signals in MitoT-red-EV treated BECs are not associated with any free MitoTracker dye co-isolated with the EVs (**Fig. 3a** and **b**).

Overall, our flow cytometry and microscopy studies results indicate that the co-isolates from the blank medium control did not show any significant signals, confirming that the observed effects in our experiments are due to the mitochondria-containing EVs and not due to any potential co-isolates.

### 3.4. Mouse BEC-20K EVs outperformed human BEC-20K EVs in increasing recipient ischemic mouse and human BEC ATP levels

We compared the effects of mBEC and hBEC-derived EVs on the resulting ATP levels in recipient OGD BECs (**Fig. 5**). We determined whether homotypic EVs derived from the same species as the recipient cell (**Fig. 5a,d**) may show higher functional efficiency than heterotypic pairs: cross-species EVs and recipient cells (**Fig. 5b,c**). Recipient mBECs and hBECs were incubated in OGD medium in a hypoxic chamber for four hours. OGD-exposed BECs were treated with hBEC- and mBEC-EVs at different concentrations for 24 h. Post-exposure, recipient BEC relative ATP levels were measured using a Cell TiterGlo-based luminescent ATP assay. The ATP levels of cells treated with complete growth medium in a humidified incubator were used as normoxic control. mBECs and hBECs incubated in OGD medium for 24 h were used as OGD control (**Fig. 5a-d**).

**Figure 5:**
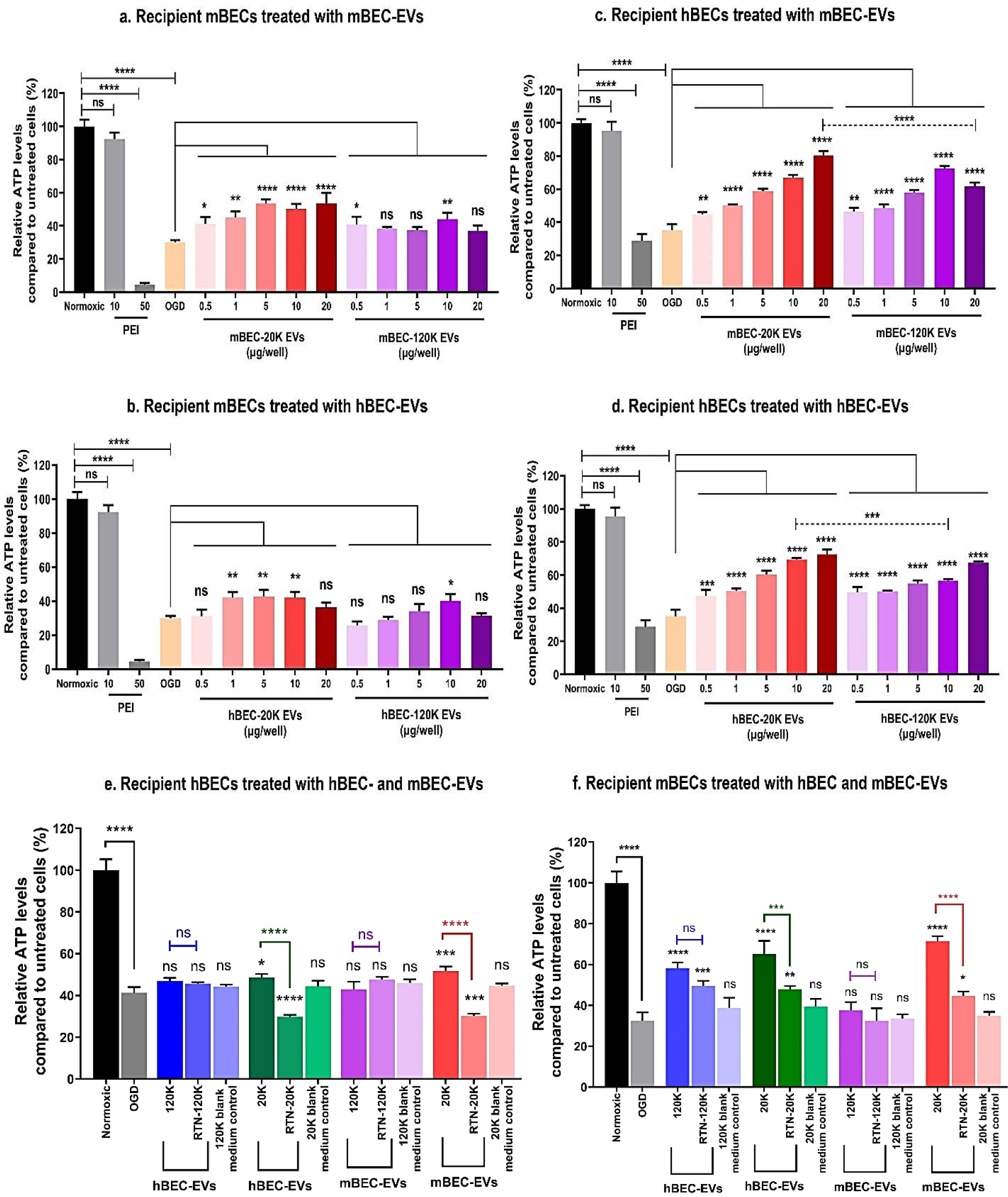
Effect of mBEC- and hBEC-EVs on recipient oxygen-glucose-deprived BEC ATP levels. mBECs and hBECs were cultured in 96-well plates in a humidified incubator at 37°C. Confluent monolayers were exposed to OGD medium in a hypoxic chamber for four hours. Post-OGD, the cell were treated with mBEC- and hBEC-120K EVs and 20K EVs at indicated amounts in OGD medium for 24 h. Cell treated with complete growth medium was used as normoxic control, whereas cells treated with only OGD medium was used as OGD control. Post-treatment, treatment mixtures were removed, and cells were incubated with a 1:1 mixture of fresh growth medium and Cell titer Glo reagent. The relative luminescence units (**RLU**) in each well were measured using a SYNERGY HTX multimode plate reader at 1 s integration time. Relative ATP levels were calculated by normalizing the RLU of treatment groups to the RLU of normoxic untreated cells in complete growth medium. The effects of mBEC-derived EVs on recipient mBECs **(a)**, hBEC-derived EVs on recipient mBECs **(b)**, mBEC-derived EVs on recipient hBECs **(c)**, and hBEC-derived EVs on recipient hBEC ATP levels **(d)** were statistically evaluated compared to OGD control. **(e,f)** *Effect of mitochondria-lacking 20K EVs on recipient hBEC and mBEC ATP levels.* OGD-exposed hBECs and mBECs were treated with mBEC and hBEC-naïve EVs, RTN-EVs, and blank medium controls at 10 μg of EV per well for 24 h in OGD conditions. RTN-EVs were isolated from donor BECs pre-exposed to 0.25 µM RTN. After incubation, the treatment mixture was removed, and the ATP assay was performed. Relative ATP levels were calculated by normalizing the RLU of treatment groups to the RLU of normoxic control. Data represents mean ± SD (n = 3). * p < 0.05, ** p < 0.01, *** p < 0.001, **** p < 0.0001, ns: non-significant.

OGD exposure led to about a 70% reduction in mBEC ATP levels compared to normoxic mBECs (**Fig. 5 a-b**), whereas OGD reduced hBEC ATP levels to about 65% (**Fig. 5c-d**). OGD-mediated decreases in BEC ATP levels were statistically significant (p<0.0001) compared to untreated normoxic cells. Recipient OGD mBECs treated with mBEC-(**Fig. 5a**) and hBEC-20K EVs (**Fig. 5b**) at a dose as low as 1 μg per well showed a statistically significant (p<0.05) increase in cellular ATP levels compared to untreated OGD mBECs. Increasing the 20K EV concentration showed a significant (p<0.0001) increase in BEC ATP levels (**Fig. 5a-b**). Notably, mBEC-20K EVs showed a greater increase in recipient mBEC ATP levels (**Fig. 5a**) compared to hBEC-20K EVs (**Fig. 5b**) at all tested doses. In contrast, OGD mBECs treated with 120K EVs showed no significant increase in cellular ATP levels (**Fig. 5a-b**), likely due to the lower mitochondrial load in 120K EVs.

Moreover, recipient OGD hBECs treated with mBEC-(**Fig. 5c**) and hBEC-20K EVs (**Fig. 5d**) showed a concentration-dependent significant (p<0.0001) increase in cellular ATP levels compared to untreated OGD hBECs. OGD hBECs treated with 120K EVs showed a statistically significant (p<0.0001) increase in cellular ATP levels (**Fig. 5c-d**). It should be noted that 20K EV-mediated increase in hBEC ATP levels was significantly (p<0.0001) greater than 120K EVs (**Fig. 5c-d**). Notably, mBEC-20K EVs increased cellular ATP levels to a greater magnitude than hBEC-20K EVs in recipient mBECs (**Fig. 5a-b**) and hBECs (**Fig. 5c-d**). In conclusion, mBEC-20K EVs outperformed hBEC-20K EVs in increasing ischemic BEC ATP levels.

Furthermore, to determine whether EV-mediated increases in recipient BEC ATP levels is a function of EV mitochondrial components, we studied the effect of EVs isolated from cells with impaired mitochondrial function. EVs with impaired mitochondrial function were isolated from the conditioned medium of hBECs or mBECs pretreated with 0.25 µM RTN. Recipient hBECs were treated with naïve 120K EVs, naïve 20K EVs, RTN-120K EVs, RTN-20K EVs, 20K- and 120K-blank medium controls for 24 h under OGD conditions. OGD hBECs (**Fig. 5e**) showed a significant (p<0.0001) decrease in cellular ATP levels compared to normoxic BEC ATP levels. OGD hBECs treated with hBEC- and mBEC-120K EVs did not show changes in ATP levels compared to untreated OGD cells (**Fig. 5e**). OGD BECs treated with hBEC- and mBEC-20K EVs showed a significant (p<0.01) increase in OGD BEC ATP levels (**Fig. 5e**). There was no significant difference between hBEC ATP levels treated with naïve 120K vs. RTN-120K EVs (**Fig. 5e**), suggesting that inhibition of donor hBEC mitochondrial function did not affect the ATP-increasing functionality of 120K EVs. Notably, RTN-20K EVs significantly reduced BECs ATP levels compared to naïve 20K EVs-treated BECs (**Fig. 5e**), suggesting that inhibition of donor cell mitochondria complex I reduced 20K EV-mediated increase in BEC ATP levels. RTN-20K EV-mediated decrease in BEC ATP levels was consistent for 20K EVs isolated from hBECs and mBECs (**Fig. 5e**). Importantly, blank medium controls did not show statistically significant changes in BEC ATP levels compared to untreated OGD BECs. This indicates that RTN-20K EV-mediated decrease in cellular ATP levels is likely due to impaired mitochondria in RTN-20K EVs, rather than a bias effect of RTN co-isolates from the conditioned medium during sequential centrifugation. Overall, our results suggested that the 20K EV-mediated increase in cellular ATP levels, regardless of EV donor species, is a function of mitochondria in 20K EVs. The suppression of relative ATP levels was much more profound in the 20K EVs than in the 120K EVs, suggesting that the increase in ATP mediated by 20K EVs is even more dependent on mitochondrial complex I function.

A similar trend was observed in experiments involving recipient mBECs. OGD mBECs showed a significant decrease in ATP levels, with 20K EVs from hBECs and mBECs significantly increasing ATP levels (p<0.001, **Fig. 5f**). In contrast, mBEC-derived 120K EVs did not increase ATP levels (**Fig. 5f**). RTN-20K EVs significantly reduced ATP levels, indicating the crucial role of functional mitochondria in the 20K EV-mediated ATP increases. Additionally, RTN-medium controls did not show changes in mBEC ATP levels (relative to untreated OGD cells) (**Fig. 5f**), suggesting that the RTN-20K EV-mediated decrease is due to impaired activity of RTN-20K EVs and not due to RTN co-isolates.

### 3.5. mBEC-20K EVs outperformed hBEC-20K EVs in increasing mitochondrial respiration of recipient OGD mBECs and showed anti-ROS effects

The effect of mBEC- or hBEC-EV exposure on recipient mBEC mitochondrial function was evaluated using the Seahorse analysis (**Fig. 6a**). OGD BECs were treated with different doses of EVs diluted in OGD medium for 24 h. The selected doses: 0.3, 3.3 and 13.2 µg EV protein are equivalent to ∼3, 30 and 120 µg/cm^2^ per well of the Seahorse plate (surface area per well: 0.11 cm^2^). First, recipient mBECs exposed to OGD conditions significantly decreased OCR compared to untreated normoxic control (**Fig. 6a**), suggesting that OGD significantly reduced cellular oxidative phosphorylation. OGD mBECs treated with mBEC- and hBEC-120K EVs did not show any statistical increase in oxygen consumption rate (**OCR**) compared to untreated OGD control (**Fig. 6a**). A low dose of 20K EVs (0.3 μg/well) did not increase mitochondrial respiration of OGD mBECs (**Fig. 6a**). mBECs treated with 3.3 and 13.3 μg/well mBEC- and hBEC-20K EVs showed a statistically significant (mBEC-20K EVs: p<0.001, hBEC-20K EVs: p<0.05) increases in OCR compared to untreated OGD control (**Fig. 6a**). Notably, mBEC-20K EVs increased mBEC mitochondrial respiration to significantly (p<0.001) greater extent than hBEC-20K EVs (**Fig. 6a**). Overall, hBEC- and mBEC-20K EVs significantly increased recipient mBEC and hBEC mitochondrial respiration compared to untreated and 120K EV-treated ischemic mBECs. Homotypic mBEC-20K EVs outperformed hBEC-20K EVs in increasing recipient mBECs mitochondrial respiration. Collectively, our data suggests that 20K EVs derived from the same species as the recipient species, a.k.a., homotypic pairs (recipient mBECs treated with mBEC-20K EVs), show improved mitochondrial respiration than heterotypic pairs.

**Figure 6:**
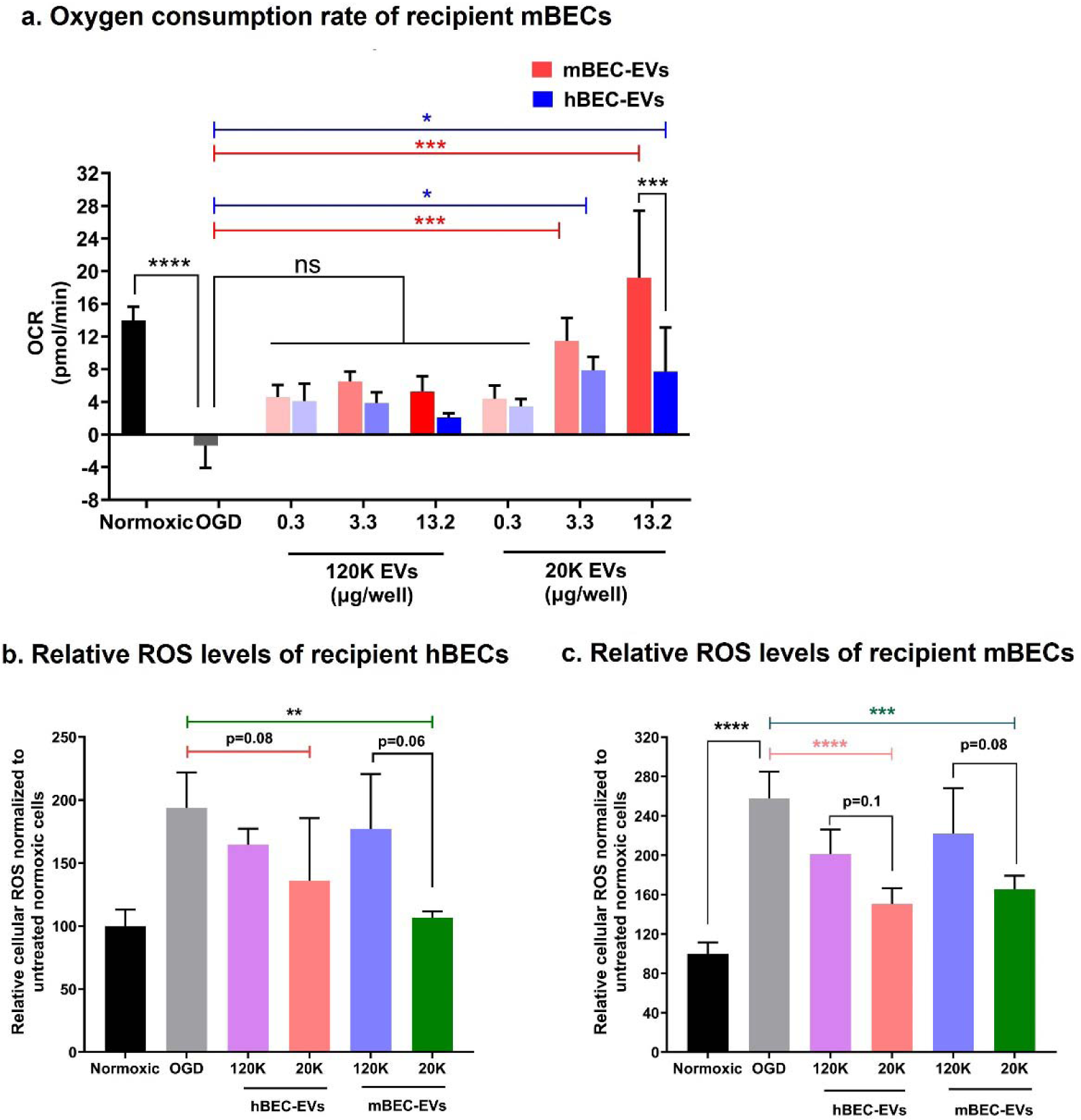
Effect of mouse and human BEC-derived EVs on recipient mBEC mitochondrial respiration and ROS levels under OGD conditions. ***(a) Effect on recipient OGD mBEC OCR levels***: mBECs were cultured in a Seahorse XF96 plate for four days at 20,000 cells/well. Confluent mBECs were incubated with OGD medium in a hypoxic chamber for four hours. Post-OGD exposure, cells were incubated with mBEC and hBEC-EVs at indicated doses diluted in OGD medium for 24 h. Post-treatment, the medium was replaced with DMEM. Maximum oxygen consumption rate (OCR) in mBEC was analyzed using the Seahorse XFe96 analyzer. Data represents mean±SD (n=3). * p<0.05, ** p<0.01, *** p<0.001, **** p<0.0001. *(b,c) Anti-ROS effect of hBEC and mBEC-derived EVs under OGD conditions:* Recipient hBECs **(b)** and mBECs **(c)** were treated with hBEC- or mBEC-derived 120K and 20K-EVs at 10 μg EV protein/well for 24 h. Untreated cells under OGD conditions and untreated, normoxic cells incubated in complete growth medium were used as OGD and normoxic controls, respectively. Post-treatment cells were treated with CellROX deep red reagent for 30 min and washed thrice with PBS. Relative fluorescence intensities were measured at excitation and emission wavelengths of 640 nm and 665 nm, respectively, using a Tecan Infinite M1000 plate reader. Relative ROS levels were calculated by normalizing the fluorescence intensities of treated groups to those of normoxic control, untreated cells.

We performed additional studies to determine EV-mediated anti-ROS effects of recipient BECs under OGD conditions (**Fig. 6b and c**). CellROX oxidative stress reagent was utilized to evaluate the effect of OGD on cellular ROS levels in EV-treated recipient BECs. Recipient hBECs exposed to OGD conditions showed a statistically significant (p<0.0001) increase in hBEC ROS levels (**Fig. 6b**). OGD hBECs treated with hBEC- and mBEC-derived 120K EVs did not show any reduction in cellular ROS levels, whereas 20K EVs showed a considerable decrease in hBEC ROS levels (**Fig. 6b**). Similarly, mBECs exposed to OGD showed 2.5-fold and a statistically significant (p<0.0001) increase in mBEC ROS levels (**Fig. 6c**). OGD mBECs treated with hBEC- and mBEC-derived 120K EVs did not show any significant decrease in cellular ROS levels (**Fig. 6c**). Notably, recipient OGD mBECs treated with hBEC- and mBEC-derived 20K EVs showed a statistically significant (p<0.001) reduction in mBEC ROS levels compared to untreated OGD mBECs (**Fig. 6c**). The 20K EV-mediated reductions in ROS levels showed a trend albeit with p>0.05 and with ROS levels comparable to control, untreated normoxic cells (**Fig. 6c**). There was no significant difference in anti-ROS effects between human and mouse BEC-derived 20K EVs (**Fig. 6c**). In conclusion, BEC-derived 20K EVs showed a trend towards anti-ROS effects in OGD BECs and to a greater magnitude compared to 120K EVs.

### 3.6. Intravenously injected mBEC-20K EVs demonstrate improved post-stroke outcomes in a mouse middle cerebral artery occlusion (MCAo) model of ischemic stroke

We evaluated the therapeutic efficacy of mBEC-20K EVs on resulting mouse brain infarct volumes and neurological functions in a mouse MCAo model of ischemic stroke. The mouse MCA was occluded using a silicon filament for 90 min to induce cerebral ischemia in mice, and reperfusion was initiated by withdrawing the filament. mBEC-derived 20K EVs at 100 μg total EV protein in 200 μL of PBS were intravenously injected two hours after ischemic onset, and mice treated with PBS were used as vehicle control. Twenty-four hours post-treatment, mice were euthanized, and brain infarct volumes were analyzed using TTC-stained sections by a blinded investigator. The brain sections of mice treated with mBEC-20K EVs showed notably reduced infarct volume (white region) and showed greater viable tissue (red region) compared to the infarct volumes in mice treated with vehicle control (**Fig. 7a**). A quantitative analysis of the total hemispheric infarct volume showed that 20K EV-treated mice brain sections showed about a 45% reduction (p<0.01) in infarct volume compared to PBS-treated mice (**Fig. 7b**).

**Figure 7:**
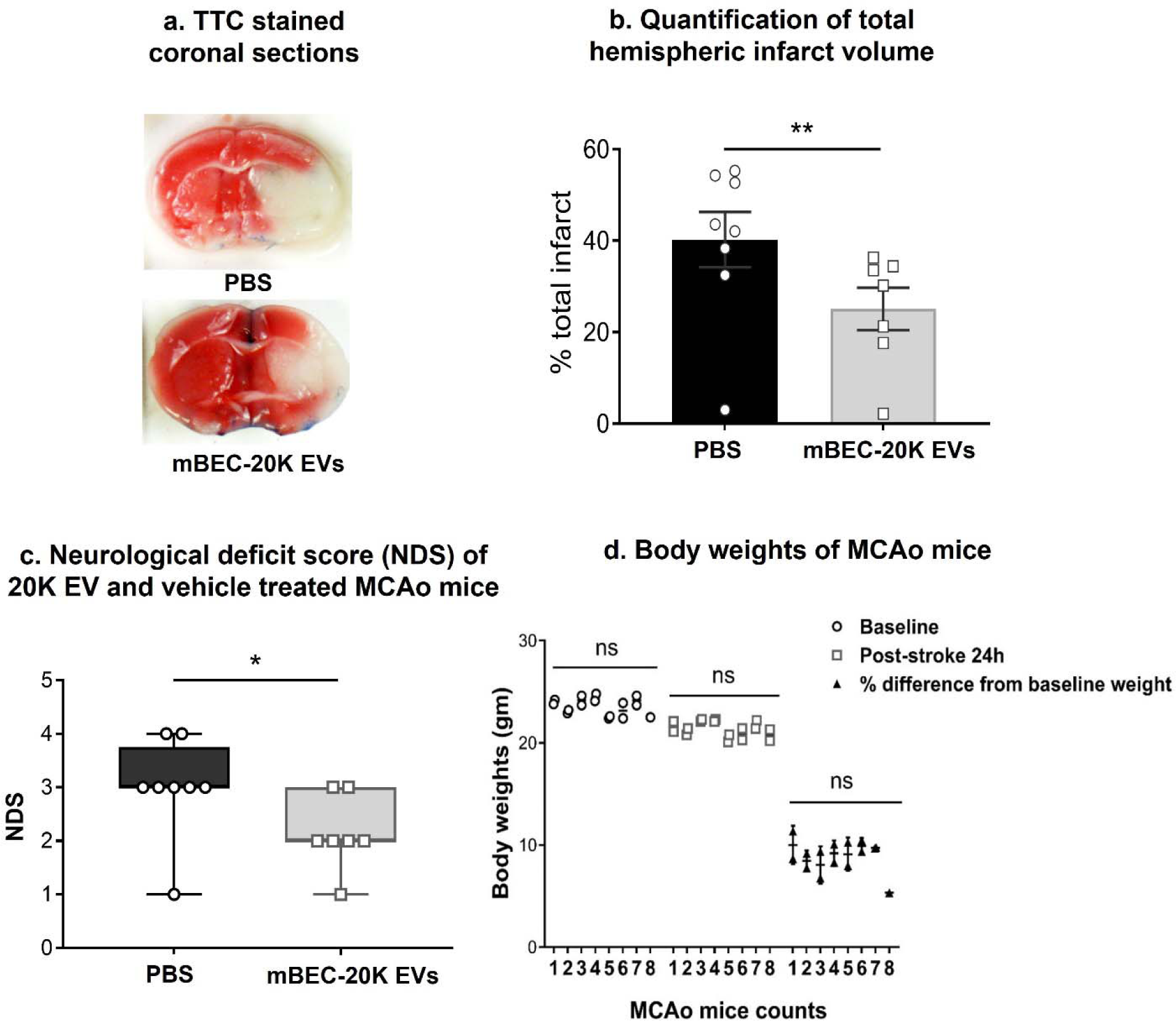
*IV*-administration of mBEC-derived mitochondria-containing 20K EVs showed neuroprotection in a mouse MCAo model of stroke. Young male C57BL/6 mice (8–12 weeks) were subjected to middle cerebral artery occlusion for 90 min to induce ischemic stroke. Reperfusion was initiated by withdrawing the filament. Two hours after ischemic onset, mice were treated with either mBEC-20K EVs (100 μg EV protein in 200 μL PBS) or vehicle control (200 μL of PBS). Twenty-four hours post-reperfusion, mice were euthanized and brains were analyzed for infarct size using TTC-stained sections. **(a)** representative TTC stained coronal section of PBS- or 20K EV-treated MCAo mice. **(b)** Quantitative analysis of total hemispheric brain section infarct volume of 20K EVs (n=7) or PBS (n=8) treated mice. The data represents mean±SEM. The statistical difference between treatment and control group was performed using unpaired student’s t-test. **(c)** Behavioral outcomes were measured using NDS at 24 hours post-treatment in 20K EV- and PBS-treated MCAo mice. The statistical difference between treatment and control group was determined using the Mann-Whitney U test. *p<0.05, **p<0.01. **(d)** Body weights of PBS and mBEC-20K EV-treated MCAo mice at baseline and 24 h post-stroke to determine the gross safety of mBEC-20K EV therapy. Body weights of MCAo mice in vehicle and EV-treated groups were evaluated at stroke onset (baseline) and 24 h post-treatment.

We further studied the effects of 20K EV administration on behavioral outcomes in the MCAo mice, again using a blinded investigator (**Fig. 7c**). Twenty-four hours post-treatment, mice were observed for their locomotor activities, and neurological deficit scores (**NDS**) of 20K EV-treated mice were compared against vehicle controls. NDS scores of 20K EV-treated mice were significantly (p<0.05) lower compared to vehicle controls, suggesting that *IV*-administration of mBEC-derived mitochondria-containing 20K EVs significantly improved post-stroke behavioural outcomes of MCAo mice.

We also did not observe any difference in treatment-related mortalities, suggesting that mBEC-20K EVs are safe when administered intravenously to the mice. Additionally, to determine the safety of mBEC-20K EV therapy, body weights of MCAo mice in vehicle and EV-treated groups were evaluated at stroke onset (baseline) and 24 h post-treatment. The average body weight of vehicle mice at baseline and 24 h post-stroke were 23.6 and 21.5 gm, respectively (**Fig. 7d**). Moreover, the average body weight of mBEC-20K EV treated mice at baseline and 24 h post-stroke were 23.5 and 20.2 gm, respectively (**Fig. 7d**). The average body weight loss from baseline for vehicle and EV-treated groups was 8.8% and 9.1%, respectively (**Fig. 7d**). Hence, there is no significant difference in post-stroke weight loss between EV and vehicle-treated groups at 24h after stroke, demonstrating the safety of *IV*-administered mBEC-20K EVs.

We investigated the therapeutic efficacy of hBEC-derived 20K EVs on mouse MCAo brain infarct volumes and previously published representative 2,3,5-triphenyl tetrazolium chloride (**TTC**) -stained sections and infarct volume data [33]. Briefly, in that pilot experiment, MCAo mice were treated with 100 μg of hBEC-20K EVs or a vehicle control two hours after stroke onset. Mice were euthanized 24 hours post-stroke, and their brains were analyzed for infarct size using TTC staining (**SL Fig. 7a**). The representative TTC image showed that MCAo mice treated with hBEC-20K EVs showed a greater fraction of viable ipsilateral brain tissue compared to PBS-treated group. Quantitative analysis showed that hBEC-20K EV treated mice showed about a 35% reduction in total hemispheric infarct volume compared to PBS-treated control MCAo mice. The decrease in infarct volume was not statistically significant; however, it showed a trend towards neuroprotection in a small cohort of the 20K EV-treated mice compared to the vehicle-treated group (**SL Fig. 7**).

## 4. Discussion

This study investigated the effect of homotypic *vs*. heterotypic EV donor cell-recipient cell pair on 20K EV mitochondria functionality in the recipient BECs. We evaluated the effect of mBEC-derived mitochondria-containing 20K EVs on the metabolic function of recipient ischemic BECs and neuroprotective effects in a mouse MCAo model of stroke. Ischemic stroke-induced OGD leads to mitochondrial dysfunction in BECs, resulting in cell death and BBB breakdown. We rationalized that delivery of exogenous 20K EV mitochondria into BECs can restore cellular metabolic functions and limit post-stroke neurological dysfunction [35].

EV donor cell type and species are factors that can significantly contribute to the therapeutic efficiencies of EVs [35, 36]. BECs contain about five-fold greater amounts of mitochondria than peripheral endothelial cells [8, 9]; and therefore, BEC-derived EVs potentially allow incorporation of a greater innate mitochondrial load than EVs from other cell types. BEC-EV membranes contain surface proteins such as transferrin receptor, insulin receptor, integrins, intercellular adhesion molecules, and CD46 [45–48], facilitating EV interactions with BBB surface receptors and subsequently allowing internalization into recipient BECs. Our previous study demonstrated that hBEC-20K EVs contain mitochondria [33, 38]. hBEC-20K EVs transferred mitochondria that colocalized with the mitochondrial network of the recipient BECs. As a consequence, hBEC-20K EVs increased ischemic hBEC ATP levels and mitochondrial respiration. A pilot study (n=4 mice) of *IV* injected hBEC-20K EVs in a mouse MCAo model of ischemic stroke showed a trend towards a potential efficacy signal [33]. We rationalized that a heterotypic combination of EV source species (hBEC-EVs) and recipient animal species (mouse model of stroke) likely contributed to the sub-optimal therapeutic efficacy of hBEC-20K EVs in the mouse stroke model. Therefore, in this work, we hypothesized that *IV* administration of *m*BEC-20K EVs may show a superior reduction in brain infarct volume and improve neurological functions in MCAo mice.

In this work, we compared mBEC-*vs*. hBEC-20K EVs in terms of their (1) physicochemical properties and biomarkers and (2) capabilities to transfer the innate 20K EV mitochondrial load into recipient mBECs or hBECs—we compared the levels of transfer between homotypic *vs*. heterotypic EV donor cell-recipient cell pairs. We further determined if ischemic conditions can affect 20K EV mitochondria transfer into recipient OGD BECs. Finally, we evaluated the therapeutic efficacy of mBEC-20K EV in a mouse MCAo model of ischemic stroke. Our results showed that irrespective of the donor cell species, both mBEC- and hBEC-20K EVs contained mitochondria, and 20K EVs lacked mitochondria. mBEC-EVs transferred their mitochondrial components into both recipient mBECs and hBECs. OGD conditions significantly reduced 20K EV mitochondrial transfer compared to normoxic conditions. mBEC-20K EVs outperformed hBEC-20K EVs in increasing recipient ischemic mBEC ATP levels and oxygen consumption rate. Rotenone-mediated inhibition of donor BEC mitochondrial function significantly reduced 20K EV mitochondria ATP-increasing functionality in recipient BECs, confirming that 20K EV-mediated increases in recipient ischemic BEC ATP levels were a function of innate 20K EV mitochondria. Lastly, *IV*-injected mBEC-20K EVs showed a statistically significant reduction in mouse brain infarct volume and significantly improved neurological functions in a mouse MCAo model of ischemic stroke.

We isolated 120K EVs and 20K EVs from the conditioned medium of mBEC and hBECs using a differential ultracentrifugation method— a commonly used EV isolation method [51, 52]. 120K EVs showed characteristic particle diameters <200 nm, whereas the diameters of 20K EVs were >200 nm (**Fig. 1a**), aligning with previous reports [18, 25, 34, 37, 53]. mBEC-20K EVs showed significantly higher particle diameters than hBEC-20K EVs (**Fig. 1a**). The net negative zeta potentials on EV membrane (**Fig. 1a**) are likely due to EV membrane phospholipid components such as phosphatidylinositol, phosphatidylserine, and glycosylated lipid derivatives [54, 55]. The natural process of EV biogenesis likely results in heterogenous EV sub-populations, manifesting as broad dispersity indices (**Fig. 1a**) that seemed consistent for human and mouse species-derived EVs. We observed similarities in the morphology of 120K EVs and 20K EVs irrespective of EV donor species (mouse *vs*. human, **Fig. 1b**). It is important to note that 20K EVs are a heterogeneous subpopulation with dispersity indices ranging from 0.3 to 0.5 (**Fig. 1a**). The particle size distribution of 20K EVs shows that about 75% of the measured sample ranges from 150 to 850 nm (**SL Fig. 3**). **Fig. 1a** reports intensity-weighted z-average particle diameters using DLS, while **Fig. 1b** presents representative TEM images of negatively-stained EVs. The differences in observed particle sizes between DLS and TEM are due to their distinct measurement principles and sample handling conditions. DLS estimates the hydrodynamic diameter, which includes the particle core and the surrounding liquid layer in suspension. In contrast, TEM measures the core size of individual particles in a dried state under vacuum, potentially shrinking the particles due to solvent diffusion. Moreover, TEM represents only a small portion of the sample, which can be influenced by operator bias and is limited by the number of images and particles analyzed. Accurate analysis of heterogeneous samples like EVs requires multiple images to determine the true particle size distribution of the bulk sample [56].

EV characterization using western blotting (**Fig. 1c**) showed that 120K EVs and 20K EVs have minimal endoplasmic reticulum contaminants, suggesting the purity of isolated EVs. CD63 and CD9, tetraspanin markers associated with exosomal cargo selection, binding, and uptake of 120K EVs by target cells [57], were expressed in 120K EVs derived from both species. ADP-ribosomal factor, **ARF6**, a GTP binding protein potential for cell migration/invasion, was expressed in 20K EVs, consistent with a published report [58]. Collectively, human and mouse BEC-derived EVs expressed characteristic protein biomarkers.

TEM analysis of cross-sectioned EVs showed that 20K EV contained one or multiple mitochondria with heterogenous sizes and shapes in the 20K EV lumen, whereas 120K EVs lack electron-dense structures (**Fig. 2a**). Notably, the mitochondrial structures in 20K EVs (**Fig. 2a**) were similar to the extracellular free mitochondria isolates and mitochondria in EV lumen reported in the published literature [18, 19, 50, 59]. Phinney *et al.* demonstrated the selective packaging of mitochondria in 20K EVs while 120K EVs carried mtDNA, but not mitochondria [18]. The authors demonstrated that overexpression of Phosphatase and tensin homolog-induced kinase 1/Parkin proteins and suppression of Miro 1/2 protein on the outer mitochondrial membrane leads to mitochondria-cytoskeletal interactions [18]. The resulting complexes migrate towards the plasma membrane, package into cell buds, and pinching of these cell buds secretes mitochondria-containing 20K EVs into extracellular spaces [18]. We further confirmed the selective presence of mitochondrial structural protein TOMM20 (**Fig. 2b**), an outer mitochondrial membrane protein that regulates the import of specific proteins from the cytosol [60], confirming the presence of mitochondria in 20K EVs but not in 120K EVs aligning with results of our TEM analysis (**Fig. 2a**). Overall, our results aligned with published reports [19, 32] demonstrating the presence of mitochondrial proteins in 20K EVs. Ikeda *et al.* showed the presence of ATP5A proteins in human cardiomyocyte-derived mitochondria-containing 20K EVs [19], whereas Silva *et al.* showed human mesenchymal stromal cell-derived 20K EVs expressed TOMM20 [32].

We further investigated the effects of (1) donor EV and recipient BEC species and (2) oxygen-glucose deprived *vs*. normoxic conditions on the extent of EV mitochondrial transfer. Interestingly, mBEC-20K EVs showed a greater extent of mitochondria transfer into recipient normoxic BECs than mBEC-120K EVs, hBEC-20K EVs and hBEC-120K EVs at the highest tested dose (**Fig. 3c-d**). Noteworthy, mBEC-20K EVs showed substantially higher average 20K EV particle diameters than hBEC-20K EVs (**Fig. 1a**). We speculate that mBECs produce large 20K EVs with a greater mitochondrial load than hBEC counterparts. As a result, at the same total EV protein dose, mBEC-20K EVs showed greater mitochondrial transfer into recipient BECs than hBEC-20K EVs. Our results align with the *in vivo* study by Banks *et al.* [20] that investigated the effect of parent cell line species using EVs from cancerous *vs*. non-cancerous cells on EXOs (120K EVs) pharmacokinetics, brain distribution, and extent of BBB crossing in mice. The authors showed that all EXOs derived from mouse, human, cancerous, and non-cancerous donor cell lines crossed BBB, but their transport from blood to brain parenchyma varied from 58% to 92% compared to albumin control [20]. Importantly, mouse cell line-derived EXOs showed greater brain transport than human cell line-derived EXOs [20]. EXO membrane proteins, including CD46, ICAM1, and integrin, facilitate their transport to the BBB via adsorptive transcytosis [20].

Furthermore, we investigated the effect of OGD conditions on 20K EV mitochondria transfer in recipient BECs using flow cytometry analysis. Alteplase-mediated dissolution of blood clots, the front-line treatment for stroke, restores blood flow (*i.e.,* reperfusion). Our goal is to deliver mitochondria-containing 20K EVs during the reperfusion phase but OGD-mediated reduction in BEC metabolic functions may affect their EV uptake efficiency compared to healthy (normoxic) conditions. Therefore, we wanted to determine if OGD BECs can also internalize BEC-EVs. Our results demonstrated MitoT+ve cells in OGD BECs in all homotypic *vs*. heterotypic EV donor-recipient cell pairs (**Fig. 4 a-d**), indicating the potential of 20K EVs to be internalized during ischemic conditions as well as normoxic conditions. OGD conditions likely compromised cellular metabolic functions, translating to reduced EV internalization into recipient BECs. Hema *et al.* showed that prolonged hypoxic exposure decreased clathrin-independent endocytic uptake while clathrin-mediated endocytosis remained unaffected [61].

Mitochondrial oxidative phosphorylation produces 80-90% of cellular ATP, and OGD-induced mitochondrial dysfunction significantly reduces cellular ATP levels. Our data showed that four hours of OGD exposure reduces about 60% of BEC ATP levels compared to untreated normoxic BECs, which is consistent with the published reports [11, 62]. Our data showed that mBEC-20K EVs outperformed in increasing recipient OGD BEC ATP levels compared to mBEC-120K EVs, hBEC-120K EVs, and hBEC-20K EVs. Importantly, rotenone-mediated inhibition of donor BEC mitochondrial function significantly reduced ATP-increasing functions of 20K EVs to a much greater extent than RTN-120K EVs. This data suggested that an 20K EVs -mediated increase in recipient BEC ATP levels is a function of 20K EV mitochondria, which is in agreement with our previous studies [33, 38]. As 120K EVs lack mitochondria, rotenone-mediated inhibition of donor mitochondria minimally affected 120K EV-mediated increases in ATP levels.

O’Brien *et al.* demonstrated that human mesenchymal stromal cell-derived 20K EVs increased recipient human cardiomyocyte ATP levels [49]. The authors showed that 20K EV transferred their mitochondria into recipient cells and colocalized with endogenous cell mitochondria. 20K EV treatment increased cellular ATP levels and suppressed apoptosis in doxorubicin-injured cardiomyocytes [49]. Silva *et al.* showed that human pulmonary microvascular endothelial cells treated with human MSC-derived 20K EVs showed a three-fold increase in recipient cellular ATP levels and a six-fold increase in mitochondrial respiration [32]. Rhodamine-mediated inhibition of 20K EV mitochondrial functions did not increase endothelial ATP levels and mitochondrial respiration, suggesting that 20K EV-mediated increase in recipient cell bioenergetics is a function of the innate 20K EV mitochondrial load [32].

We further evaluated the homotypic *vs.* heterotypic EV donor-recipient cell effect on the EV-mediated modulation in recipient ischemic mBEC mitochondrial respiration using Seahorse assays. The Seahorse extracellular flux analyzer facilitates real-time measurements of the oxygen consumption rate, an indicator of mitochondrial respiration, and extracellular acidification rate, an indicator of glycolytic capacity in live intact cells [63, 64]. We observed that irrespective of the EV donor BEC species, 120K EVs did not increase OGD BEC mitochondrial respiration (**Fig. 6)**. mBEC-20K EVs increased cellular oxidative phosphorylation compared to hBEC-20K EVs in the recipient mBECs (**Fig. 6**). Our data align with the published reports [18, 19, 32, 34] on EV-mediated increase in recipient cellular mitochondrial respiration. For instance, Ikeda *et al.* showed human cardiomyocyte-derived mitochondria-rich 20K EVs treated with hypoxic human cardiomyocytes showed a significant increase in basal, ATP-induced, and maximum mitochondrial respiration compared to untreated hypoxic cells [19]. Our results indicate that homotypic donor EV-recipient cell pairs outperform heterotypic donor EV-recipient cells in increasing recipient BEC mitochondrial functions.

Collectively, our data thus far showed that mBEC-20K EVs outperformed hBEC-20K EVs in 20K EV mitochondrial transfer (**Fig. 3 and 4**), increasing ischemic BEC ATP levels (**Fig. 5**), and mitochondrial respiration (**Fig. 6**). Based on the superior performance of mBEC-20K EVs, we *IV*-administered mBEC-20K EVs after ischemia/reperfusion injury in a mouse MCAo model of ischemic stroke. Our data revealed a statistically significant reduction in brain infarct volume compared to vehicle-treated mice (**Fig. 7a-b**). Importantly, mBEC-20K EVs were administered two hours after reperfusion, simulating a clinical scenario of delayed administration, and yet, we noted a 45% reduction in infarct volume to vehicle-injected mice. The therapeutic effect of IV-injected mBEC-derived 20K EVs was determined by measuring the infarct volume in brain sections using 2,3,5-triphenyltetrazonium chloride (**TTC**) method. TTC is a water-soluble and colorless dye, which is reduced by mitochondrial enzyme succinate dehydrogenase of living cells and converted into a water-insoluble light-sensitive compound: formazan. Healthy brain sections submerged in TTC solution shows deep red tissue staining. In contrast, damaged/dead tissue remains white, showing the absence of living cells, and indicating an infarcted region. As TTC is reduced by mitochondrial enzyme succinate dehydrogenase in live cells, resulting in deep red tissue staining, it is likely that the deep red regions in mBEC-treated brain sections may be suggestive of increased mitochondrial content as a result of mitochondria-containing 20K EV accumulation in the brain tissue. Additionally, the decrease in infarct volume was also accompanied by a reduction in neurological deficit score in mBEC-20K EV-treated mice compared to saline-treated mice, suggestive of EV mitochondria-mediated cerebroprotection. Importantly, mice treated with mBEC-20K EVs showed a statistically significant improvement (p=<0.5) in behavioral outcomes post-ischemia/reperfusion injury (**Fig. 7c**). It should be noted that this is the first demonstration of mBEC-20K EV-mediated cerebroprotection and improved behavioral outcomes in a mouse model of ischemic stroke.

It should be noted that we used young male mice for MCAo studies as the model has been standardized, allows to exclude estrogen-related neuroprotection effects, and the mortality rates are much lower in younger mice compared to aged mice. 20K EV administration did not show any treatment-related mortalities and showed significantly improved neurological functions 24 h post-stroke onset. However, we acknowledge that the mice were analyzed post a short end-point, and therefore, we intend to measure chronic post-stroke recovery (72 h post-administration of treatment or later) and in aged mice in future studies. In addition, our future studies will (1) evaluate the effects of different 20K EV doses, (2) determine whole blood and brain pharmacokinetics and systemic biodistribution of 20K EV mitochondria in healthy and MCAo mice, and (3) determine therapeutic efficacy of 20K EV mitochondria in aged male and female mice. The cell assays we have reported here in were not designed to detect any potential sex differences in EV mitochondrial load. Pulmonary ECs from male mice showed an increased basal and maximal respiration rates, and were more sensitive to pharmacologically-impaired mitochondrial function compared to cells from female mice [65]. If our future experiments reveal sex differences, we will compare primary BECs [66] isolated from male *vs*. female mice in our subsequent studies. Our future studies will evaluate the accumulation of Mitotracker-labeled 20K EV deposition in brain and brain microvessels post-IV administration. The presence of Mitotracker signals in the brain sections would determine 20K EV mitochondria transfer into the brain. The correlation between Mitotracker-20K EV accumulation and a decrease in infarct volume would establish the association of therapeutic efficacy with 20K EV mitochondria accumulation in the brain. Additionally, future studies will evaluate the mitochondrial respiration in isolated brain microvessels of mBEC-20K EV-treated MCAo mice using Seahorse analysis. Additionally, future studies will also evaluate nuclear antigen Ki-67 expression in the brain sections to determine if the increased ATP levels reflect cell proliferation and not just the transfer of mitochondrial components from 20K EVs to recipient BECs.

## 5. Conclusions

Our results suggest that mBEC-20K EVs likely contain greater mitochondrial load than hBEC-20K EVs. Homotypic donor EV-recipient cell pairs outperform heterotypic donor EV-recipient cells in increasing recipient BEC ATP levels and mitochondrial respiration. *IV* administration of mBEC-20K EVs decreased brain infarct volume and improved behavioral outcomes compared to vehicle-treated controls in a mouse model of ischemic stroke. Our data demonstrates that mBEC-derived mitochondria-containing 20K EVs is a potent approach to improve post-stroke outcomes.

## 6. Author contributions

Conceptualization, Formal analysis, Methodology, Funding acquisition, Project administration, Resources, Supervision D.S.M Investigation, Data curation, Formal analysis, Methodology K.M.D, V.R.V, D.B.S., S.S.S, L.D.M and D.S.M Transmission electron microscopy K.M.D and D.B.S *In vitro* assays K.M.D, B.B, E.E.H, D.G *In vivo* experiments V.R.V, V.A.Q, M.E.M Seahorse studies K.M.D, K.S.R, D.S.M. Physicochemical characterization K.M.D, E.E.H, D.G, M.C, J.M, S.Z Flow cytometry K.M.D, D.S.M Fluorescence microscopy K.M.D, D.S.M Data analysis K.M.D, D.S.M Manuscript writing K.M.D., V.V., L.D.M., D.S.M.

## 7. Acknowledgements

This work was supported via start-up funds for the Manickam laboratory from Duquesne University (DU) to the PI. We would like to acknowledge the Neurodegenerative Undergraduate Research Experience for funding EEH and BB through a grant from the National Institute of Neurological Disorders and Stroke (R25NS100118). The authors are thankful to Dr. Rehana K. Leak (DU) for the Olympus epifluorescence microscope and Odyssey imager support. The authors are grateful to Drs. Lauren O’Donnell, Manisha Chandwani, and Yashika Kamte (DU) for flow cytometry support.

## Data availability

The raw/processed data required to reproduce these findings can be obtained from the corresponding author upon request.

## References

1. Liu F, Lu J, Manaenko A, Tang J, Hu Q. Mitochondria in Ischemic Stroke: New Insight and Implications. Aging Dis. 2018;9(5):924–37.

2. Yang J-L, Mukda S, Chen S-D. Diverse roles of mitochondria in ischemic stroke. Redox Biol. 2018;16:263–75.

3. Kluge MA, Fetterman JL, Vita JA. Mitochondria and endothelial function. Circ Res. 2013;112(8):1171–88.

4. Bustamante-Barrientos FA, Luque-Campos N, Araya MJ, Lara-Barba E, de Solminihac J, Pradenas C, Molina L, Herrera-Luna Y, Utreras-Mendoza Y, Elizondo-Vega R, Vega-Letter AM, Luz-Crawford P. Mitochondrial dysfunction in neurodegenerative disorders: Potential therapeutic application of mitochondrial transfer to central nervous system-residing cells. Journal of Translational Medicine. 2023;21(1):613.

5. Benjamin EJ, Blaha MJ, Chiuve SE, Cushman M, Das SR, Deo R, de Ferranti SD, Floyd J, Fornage M, Gillespie C, Isasi CR, Jiménez MC, Jordan LC, Judd SE, Lackland D, Lichtman JH, Lisabeth L, Liu S, Longenecker CT, Mackey RH, Matsushita K, Mozaffarian D, Mussolino ME, Nasir K, Neumar RW, Palaniappan L, Pandey DK, Thiagarajan RR, Reeves MJ, Ritchey M, Rodriguez CJ, Roth GA, Rosamond WD, Sasson C, Towfighi A, Tsao CW, Turner MB, Virani SS, Voeks JH, Willey JZ, Wilkins JT, Wu JH, Alger HM, Wong SS, Muntner P. Heart Disease and Stroke Statistics-2017 Update: A Report From the American Heart Association. Circulation. 2017;135(10):e146–e603.

6. Abdullahi W, Tripathi D, Ronaldson PT. Blood-brain barrier dysfunction in ischemic stroke: targeting tight junctions and transporters for vascular protection. Am J Physiol Cell Physiol. 2018;315(3):C343–C56.

7. Nian K, Harding IC, Herman IM, Ebong EE. Blood-Brain Barrier Damage in Ischemic Stroke and Its Regulation by Endothelial Mechanotransduction. Frontiers in Physiology. 2020;11(1681).

8. Oldendorf WH, Cornford ME, Brown WJ. The large apparent work capability of the blood-brain barrier: A study of the mitochondrial content of capillary endothelial cells in brain and other tissues of the rat. Annals of Neurology. 1977;1(5):409–17.

9. Lee MJ, Jang Y, Han J, Kim SJ, Ju X, Lee YL, Cui J, Zhu J, Ryu MJ, Choi SY, Chung W, Heo C, Yi HS, Kim HJ, Huh YH, Chung SK, Shong M, Kweon GR, Heo JY. Endothelial-specific Crif1 deletion induces BBB maturation and disruption via the alteration of actin dynamics by impaired mitochondrial respiration. J Cereb Blood Flow Metab. 2020;40(7):1546–61.

10. Bernardo-Castro S, Sousa JA, Brás A, Cecília C, Rodrigues B, Almendra L, Machado C, Santo G, Silva F, Ferreira L, Santana I, Sargento-Freitas J. Pathophysiology of Blood–Brain Barrier Permeability Throughout the Different Stages of Ischemic Stroke and Its Implication on Hemorrhagic Transformation and Recovery. Frontiers in Neurology. 2020;11(1605).

11. Park H-H, Han M-H, Choi H, Lee YJ, Kim JM, Cheong JH, Ryu JI, Lee K-Y, Koh S-H. Mitochondria damaged by Oxygen Glucose Deprivation can be Restored through Activation of the PI3K/Akt Pathway and Inhibition of Calcium Influx by Amlodipine Camsylate. Scientific Reports. 2019;9(1):15717.

12. Tyagi N, Ovechkin AV, Lominadze D, Moshal KS, Tyagi SC. Mitochondrial mechanism of microvascular endothelial cells apoptosis in hyperhomocysteinemia. J Cell Biochem. 2006;98(5):1150–62.

13. Twig G, Elorza A, Molina AJA, Mohamed H, Wikstrom JD, Walzer G, Stiles L, Haigh SE, Katz S, Las G, Alroy J, Wu M, Py BF, Yuan J, Deeney JT, Corkey BE, Shirihai OS. Fission and selective fusion govern mitochondrial segregation and elimination by autophagy. EMBO J. 2008;27(2):433–46.

14. Manickam DS. Delivery of mitochondria via extracellular vesicles – A new horizon in drug delivery. Journal of Controlled Release. 2022;343:400–7.

15. Todkar K, Chikhi L, Desjardins V, El-Mortada F, Pépin G, Germain M. Selective packaging of mitochondrial proteins into extracellular vesicles prevents the release of mitochondrial DAMPs. Nature Communications. 2021;12(1):1971.

16. Guescini M, Genedani S, Stocchi V, Agnati LF. Astrocytes and Glioblastoma cells release exosomes carrying mtDNA. J Neural Transm (Vienna). 2010;117(1):1–4.

17. Puhm F, Afonyushkin T, Resch U, Obermayer G, Rohde M, Penz T, Schuster M, Wagner G, Rendeiro AF, Melki I, Kaun C, Wojta J, Bock C, Jilma B, Mackman N, Boilard E, Binder CJ. Mitochondria Are a Subset of Extracellular Vesicles Released by Activated Monocytes and Induce Type I IFN and TNF Responses in Endothelial Cells. Circ Res. 2019;125(1):43–52.

18. Phinney DG, Di Giuseppe M, Njah J, Sala E, Shiva S, St Croix CM, Stolz DB, Watkins SC, Di YP, Leikauf GD, Kolls J, Riches DW, Deiuliis G, Kaminski N, Boregowda SV, McKenna DH, Ortiz LA. Mesenchymal stem cells use extracellular vesicles to outsource mitophagy and shuttle microRNAs. Nat Commun. 2015;6:8472.

19. Ikeda G, Santoso MR, Tada Y, Li AM, Vaskova E, Jung J-H, O’Brien C, Egan E, Ye J, Yang PC. Mitochondria-Rich Extracellular Vesicles From Autologous Stem Cell–Derived Cardiomyocytes Restore Energetics of Ischemic Myocardium. Journal of the American College of Cardiology. 2021;77(8):1073–88.

20. Banks WA, Sharma P, Bullock KM, Hansen KM, Ludwig N, Whiteside TL. Transport of Extracellular Vesicles across the Blood-Brain Barrier: Brain Pharmacokinetics and Effects of Inflammation. Int J Mol Sci. 2020;21(12).

21. Murphy DE, de Jong OG, Brouwer M, Wood MJ, Lavieu G, Schiffelers RM, Vader P. Extracellular vesicle-based therapeutics: natural versus engineered targeting and trafficking. Experimental & Molecular Medicine. 2019;51(3):1–12.

22. Rufino-Ramos D, Albuquerque PR, Carmona V, Perfeito R, Nobre RJ, Pereira de Almeida L. Extracellular vesicles: Novel promising delivery systems for therapy of brain diseases. Journal of Controlled Release. 2017;262:247–58.

23. Liang H-B, Chen X, Zhao R, Li S-J, Huang P-S, Tang Y-H, Cui G-H, Liu J-R. Simultaneous ischemic regions targeting and BBB crossing strategy to harness extracellular vesicles for therapeutic delivery in ischemic stroke. Journal of Controlled Release. 2024;365:1037–57.

24. Haraszti RA, Didiot M-C, Sapp E, Leszyk J, Shaffer SA, Rockwell HE, Gao F, Narain NR, DiFiglia M, Kiebish MA, Aronin N, Khvorova A. High-resolution proteomic and lipidomic analysis of exosomes and microvesicles from different cell sources. Journal of extracellular vesicles. 2016;5:32570-.

25. Dozio V, Sanchez J-C. Characterisation of extracellular vesicle-subsets derived from brain endothelial cells and analysis of their protein cargo modulation after TNF exposure. Journal of Extracellular Vesicles. 2017;6(1):1302705.

26. Tricarico C, Clancy J, D’Souza-Schorey C. Biology and biogenesis of shed microvesicles. Small GTPases. 2017;8(4):220–32.

27. Théry C, Witwer KW, Aikawa E, Alcaraz MJ, Anderson JD, Andriantsitohaina R, Antoniou A, Arab T, Archer F, Atkin-Smith GK, Ayre DC, Bach JM, Bachurski D, Baharvand H, Balaj L, Baldacchino S, Bauer NN, Baxter AA, Bebawy M, Beckham C, Bedina Zavec A, Benmoussa A, Berardi AC, Bergese P, Bielska E, Blenkiron C, Bobis-Wozowicz S, Boilard E, Boireau W, Bongiovanni A, Borràs FE, Bosch S, Boulanger CM, Breakefield X, Breglio AM, Brennan M, Brigstock DR, Brisson A, Broekman ML, Bromberg JF, Bryl-Górecka P, Buch S, Buck AH, Burger D, Busatto S, Buschmann D, Bussolati B, Buzás EI, Byrd JB, Camussi G, Carter DR, Caruso S, Chamley LW, Chang YT, Chen C, Chen S, Cheng L, Chin AR, Clayton A, Clerici SP, Cocks A, Cocucci E, Coffey RJ, Cordeiro-da-Silva A, Couch Y, Coumans FA, Coyle B, Crescitelli R, Criado MF, D’Souza-Schorey C, Das S, Datta Chaudhuri A, de Candia P, De Santana EF, De Wever O, Del Portillo HA, Demaret T, Deville S, Devitt A, Dhondt B, Di Vizio D, Dieterich LC, Dolo V, Dominguez Rubio AP, Dominici M, Dourado MR, Driedonks TA, Duarte FV, Duncan HM, Eichenberger RM, Ekström K, El Andaloussi S, Elie-Caille C, Erdbrügger U, Falcón-Pérez JM, Fatima F, Fish JE, Flores-Bellver M, Försönits A, Frelet-Barrand A, Fricke F, Fuhrmann G, Gabrielsson S, Gámez-Valero A, Gardiner C, Gärtner K, Gaudin R, Gho YS, Giebel B, Gilbert C, Gimona M, Giusti I, Goberdhan DC, Görgens A, Gorski SM, Greening DW, Gross JC, Gualerzi A, Gupta GN, Gustafson D, Handberg A, Haraszti RA, Harrison P, Hegyesi H, Hendrix A, Hill AF, Hochberg FH, Hoffmann KF, Holder B, Holthofer H, Hosseinkhani B, Hu G, Huang Y, Huber V, Hunt S, Ibrahim AG, Ikezu T, Inal JM, Isin M, Ivanova A, Jackson HK, Jacobsen S, Jay SM, Jayachandran M, Jenster G, Jiang L, Johnson SM, Jones JC, Jong A, Jovanovic-Talisman T, Jung S, Kalluri R, Kano SI, Kaur S, Kawamura Y, Keller ET, Khamari D, Khomyakova E, Khvorova A, Kierulf P, Kim KP, Kislinger T, Klingeborn M, Klinke DJ, 2nd, Kornek M, Kosanović MM, Kovács Á F, Krämer-Albers EM, Krasemann S, Krause M, Kurochkin IV, Kusuma GD, Kuypers S, Laitinen S, Langevin SM, Languino LR, Lannigan J, Lässer C, Laurent LC, Lavieu G, Lázaro-Ibáñez E, Le Lay S, Lee MS, Lee YXF, Lemos DS, Lenassi M, Leszczynska A, Li IT, Liao K, Libregts SF, Ligeti E, Lim R, Lim SK, Linē A, Linnemannstöns K, Llorente A, Lombard CA, Lorenowicz MJ, Lörincz Á M, Lötvall J, Lovett J, Lowry MC, Loyer X, Lu Q, Lukomska B, Lunavat TR, Maas SL, Malhi H, Marcilla A, Mariani J, Mariscal J, Martens-Uzunova ES, Martin-Jaular L, Martinez MC, Martins VR, Mathieu M, Mathivanan S, Maugeri M, McGinnis LK, McVey MJ, Meckes DG, Jr., Meehan KL, Mertens I, Minciacchi VR, Möller A, Møller Jørgensen M, Morales-Kastresana A, Morhayim J, Mullier F, Muraca M, Musante L, Mussack V, Muth DC, Myburgh KH, Najrana T, Nawaz M, Nazarenko I, Nejsum P, Neri C, Neri T, Nieuwland R, Nimrichter L, Nolan JP, Nolte-’t Hoen EN, Noren Hooten N, O’Driscoll L, O’Grady T, O’Loghlen A, Ochiya T, Olivier M, Ortiz A, Ortiz LA, Osteikoetxea X, Østergaard O, Ostrowski M, Park J, Pegtel DM, Peinado H, Perut F, Pfaffl MW, Phinney DG, Pieters BC, Pink RC, Pisetsky DS, Pogge von Strandmann E, Polakovicova I, Poon IK, Powell BH, Prada I, Pulliam L, Quesenberry P, Radeghieri A, Raffai RL, Raimondo S, Rak J, Ramirez MI, Raposo G, Rayyan MS, Regev-Rudzki N, Ricklefs FL, Robbins PD, Roberts DD, Rodrigues SC, Rohde E, Rome S, Rouschop KM, Rughetti A, Russell AE, Saá P, Sahoo S, Salas-Huenuleo E, Sánchez C, Saugstad JA, Saul MJ, Schiffelers RM, Schneider R, Schøyen TH, Scott A, Shahaj E, Sharma S, Shatnyeva O, Shekari F, Shelke GV, Shetty AK, Shiba K, Siljander PR, Silva AM, Skowronek A, Snyder OL, 2nd, Soares RP, Sódar BW, Soekmadji C, Sotillo J, Stahl PD, Stoorvogel W, Stott SL, Strasser EF, Swift S, Tahara H, Tewari M, Timms K, Tiwari S, Tixeira R, Tkach M, Toh WS, Tomasini R, Torrecilhas AC, Tosar JP, Toxavidis V, Urbanelli L, Vader P, van Balkom BW, van der Grein SG, Van Deun J, van Herwijnen MJ, Van Keuren-Jensen K, van Niel G, van Royen ME, van Wijnen AJ, Vasconcelos MH, Vechetti IJ, Jr., Veit TD, Vella LJ, Velot É, Verweij FJ, Vestad B, Viñas JL, Visnovitz T, Vukman KV, Wahlgren J, Watson DC, Wauben MH, Weaver A, Webber JP, Weber V, Wehman AM, Weiss DJ, Welsh JA, Wendt S, Wheelock AM, Wiener Z, Witte L, Wolfram J, Xagorari A, Xander P, Xu J, Yan X, Yáñez-Mó M, Yin H, Yuana Y, Zappulli V, Zarubova J, Žėkas V, Zhang JY, Zhao Z, Zheng L, Zheutlin AR, Zickler AM, Zimmermann P, Zivkovic AM, Zocco D, Zuba-Surma EK. Minimal information for studies of extracellular vesicles 2018 (MISEV2018): a position statement of the International Society for Extracellular Vesicles and update of the MISEV2014 guidelines. J Extracell Vesicles. 2018;7(1):1535750.

28. Welsh JA, Goberdhan DCI, O’Driscoll L, Buzas EI, Blenkiron C, Bussolati B, Cai H, Di Vizio D, Driedonks TAP, Erdbrügger U, Falcon-Perez JM, Fu QL, Hill AF, Lenassi M, Lim SK, Mahoney MG, Mohanty S, Möller A, Nieuwland R, Ochiya T, Sahoo S, Torrecilhas AC, Zheng L, Zijlstra A, Abuelreich S, Bagabas R, Bergese P, Bridges EM, Brucale M, Burger D, Carney RP, Cocucci E, Crescitelli R, Hanser E, Harris AL, Haughey NJ, Hendrix A, Ivanov AR, Jovanovic-Talisman T, Kruh-Garcia NA, Ku’ulei-Lyn Faustino V, Kyburz D, Lässer C, Lennon KM, Lötvall J, Maddox AL, Martens-Uzunova ES, Mizenko RR, Newman LA, Ridolfi A, Rohde E, Rojalin T, Rowland A, Saftics A, Sandau US, Saugstad JA, Shekari F, Swift S, Ter-Ovanesyan D, Tosar JP, Useckaite Z, Valle F, Varga Z, van der Pol E, van Herwijnen MJC, Wauben MHM, Wehman AM, Williams S, Zendrini A, Zimmerman AJ, Théry C, Witwer KW. Minimal information for studies of extracellular vesicles (MISEV2023): From basic to advanced approaches. J Extracell Vesicles. 2024;13(2):e12404.

29. Islam MN, Das SR, Emin MT, Wei M, Sun L, Westphalen K, Rowlands DJ, Quadri SK, Bhattacharya S, Bhattacharya J. Mitochondrial transfer from bone-marrow–derived stromal cells to pulmonary alveoli protects against acute lung injury. Nature Medicine. 2012;18(5):759–65.

30. Hough KP, Trevor JL, Strenkowski JG, Wang Y, Chacko BK, Tousif S, Chanda D, Steele C, Antony VB, Dokland T, Ouyang X, Zhang J, Duncan SR, Thannickal VJ, Darley-Usmar VM, Deshane JS. Exosomal transfer of mitochondria from airway myeloid-derived regulatory cells to T cells. Redox Biol. 2018;18:54–64.

31. Liu D, Dong Z, Wang J, Tao Y, Sun X, Yao X. The existence and function of mitochondrial component in extracellular vesicles. Mitochondrion. 2020;54:122–7.

32. Dutra Silva J, Su Y, Calfee CS, Delucchi KL, Weiss D, McAuley DF, O’Kane C, Krasnodembskaya AD. Mesenchymal stromal cell extracellular vesicles rescue mitochondrial dysfunction and improve barrier integrity in clinically relevant models of ARDS. European Respiratory Journal. 2021;58(1):2002978.

33. Dave KM, Stolz DB, Venna VR, Quaicoe VA, Maniskas ME, Reynolds MJ, Babidhan R, Dobbins DX, Farinelli MN, Sullivan A, Bhatia TN, Yankello H, Reddy R, Bae Y, Leak RK, Shiva SS, McCullough LD, Manickam DS. Mitochondria-containing extracellular vesicles (EV) reduce mouse brain infarct sizes and EV/HSP27 protect ischemic brain endothelial cultures. J Control Release. 2023;354:368–93.

34. D’Souza A, Burch A, Dave KM, Sreeram A, Reynolds MJ, Dobbins DX, Kamte YS, Zhao W, Sabatelle C, Joy GM, Soman V, Chandran UR, Shiva SS, Quillinan N, Herson PS, Manickam DS. Microvesicles transfer mitochondria and increase mitochondrial function in brain endothelial cells. J Control Release. 2021;338:505–26.

35. Dave KM, Stolz DB, Manickam DS. Delivery of mitochondria-containing extracellular vesicles to the BBB for ischemic stroke therapy. Expert Opinion on Drug Delivery.1–20.

36. Zhu X, Badawi M, Pomeroy S, Sutaria DS, Xie Z, Baek A, Jiang J, Elgamal OA, Mo X, Perle K, Chalmers J, Schmittgen TD, Phelps MA. Comprehensive toxicity and immunogenicity studies reveal minimal effects in mice following sustained dosing of extracellular vesicles derived from HEK293T cells. J Extracell Vesicles. 2017;6(1):1324730.

37. Dave KM, Zhao W, Hoover C, D’Souza A, S Manickam D. Extracellular Vesicles Derived from a Human Brain Endothelial Cell Line Increase Cellular ATP Levels. AAPS PharmSciTech. 2021;22(1):18.

38. Dave KM, Dobbins DX, Farinelli MN, Sullivan A, Milosevic J, Stolz DB, Kim J, Zheng S, Manickam DS. Engineering Extracellular Vesicles to Modulate Their Innate Mitochondrial Load. Cell Mol Bioeng. 2022;15(5):367–89.

39. Dai X, Chen J, Xu F, Zhao J, Cai W, Sun Z, Hitchens TK, Foley LM, Leak RK, Chen J, Hu X. TGFα preserves oligodendrocyte lineage cells and improves white matter integrity after cerebral ischemia. Journal of Cerebral Blood Flow & Metabolism. 2020;40(3):639–55.

40. Cardenes N, Corey C, Geary L, Jain S, Zharikov S, Barge S, Novelli EM, Shiva S. Platelet bioenergetic screen in sickle cell patients reveals mitochondrial complex V inhibition, which contributes to platelet activation. Blood. 2014;123(18):2864–72.

41. McCullough LD, Roy-O’Reilly M, Lai Y-J, Patrizz A, Xu Y, Lee J, Holmes A, Kraushaar DC, Chauhan A, Sansing LH, Stonestreet BS, Zhu L, Kofler J, Lim Y-P, Venna VR. Exogenous inter-α inhibitor proteins prevent cell death and improve ischemic stroke outcomes in mice. The Journal of Clinical Investigation. 2021;131(17).

42. Liu F, Yuan R, Benashski SE, McCullough LD. Changes in Experimental Stroke Outcome across the Life Span. Journal of Cerebral Blood Flow & Metabolism. 2009;29(4):792–802.

43. Venna VR, Weston G, Benashski SE, Tarabishy S, Liu F, Li J, Conti LH, McCullough LD. NF-κB contributes to the detrimental effects of social isolation after experimental stroke. Acta Neuropathol. 2012;124(3):425–38.

44. McCullough LD, Zeng Z, Li H, Landree LE, McFadden J, Ronnett GV. Pharmacological inhibition of AMP-activated protein kinase provides neuroprotection in stroke. J Biol Chem. 2005;280(21):20493–502.

45. Ritzel RM, Patel AR, Spychala M, Verma R, Crapser J, Koellhoffer EC, Schrecengost A, Jellison ER, Zhu L, Venna VR, McCullough LD. Multiparity improves outcomes after cerebral ischemia in female mice despite features of increased metabovascular risk. Proc Natl Acad Sci U S A. 2017;114(28):E5673–e82.

46. Ritzel RM, Pan SJ, Verma R, Wizeman J, Crapser J, Patel AR, Lieberman R, Mohan R, McCullough LD. Early retinal inflammatory biomarkers in the middle cerebral artery occlusion model of ischemic stroke. Mol Vis. 2016;22:575–88.

47. Venna VR, Benashski SE, Chauhan A, McCullough LD. Inhibition of glycogen synthase kinase-3β enhances cognitive recovery after stroke: the role of TAK1. Learn Mem. 2015;22(7):336–43.

48. Pospichalova V, Svoboda J, Dave Z, Kotrbova A, Kaiser K, Klemova D, Ilkovics L, Hampl A, Crha I, Jandakova E, Minar L, Weinberger V, Bryja V. Simplified protocol for flow cytometry analysis of fluorescently labeled exosomes and microvesicles using dedicated flow cytometer. J Extracell Vesicles. 2015;4:25530.

49. O’Brien CG, Ozen MO, Ikeda G, Vaskova E, Jung JH, Bayardo N, Santoso MR, Shi L, Wahlquist C, Jiang Z, Jung Y, Zeng Y, Egan E, Sinclair R, Gee A, Witteles R, Mercola M, Svensson KJ, Demirci U, Yang PC. Mitochondria-Rich Extracellular Vesicles Rescue Patient-Specific Cardiomyocytes From Doxorubicin Injury: Insights Into the SENECA Trial. JACC CardioOncol. 2021;3(3):428–40.

50. Hayakawa K, Esposito E, Wang X, Terasaki Y, Liu Y, Xing C, Ji X, Lo EH. Transfer of mitochondria from astrocytes to neurons after stroke. Nature. 2016;535(7613):551-5.

51. Yang D, Zhang W, Zhang H, Zhang F, Chen L, Ma L, Larcher LM, Chen S, Liu N, Zhao Q, Tran PHL, Chen C, Veedu RN, Wang T. Progress, opportunity, and perspective on exosome isolation - efforts for efficient exosome-based theranostics. Theranostics. 2020;10(8):3684–707.

52. Livshits MA, Khomyakova E, Evtushenko EG, Lazarev VN, Kulemin NA, Semina SE, Generozov EV, Govorun VM. Isolation of exosomes by differential centrifugation: Theoretical analysis of a commonly used protocol. Scientific Reports. 2015;5(1):17319.

53. Kanada M, Bachmann MH, Hardy JW, Frimannson DO, Bronsart L, Wang A, Sylvester MD, Schmidt TL, Kaspar RL, Butte MJ, Matin AC, Contag CH. Differential fates of biomolecules delivered to target cells via extracellular vesicles. Proceedings of the National Academy of Sciences. 2015;112(12):E1433–E42.

54. Konoshenko MY, Lekchnov EA, Vlassov AV, Laktionov PP. Isolation of Extracellular Vesicles: General Methodologies and Latest Trends. Biomed Res Int. 2018;2018:8545347.

55. El Andaloussi S, Mäger I, Breakefield XO, Wood MJA. Extracellular vesicles: biology and emerging therapeutic opportunities. Nature Reviews Drug Discovery. 2013;12(5):347–57.

56. Erdbrügger U, Lannigan J. Analytical challenges of extracellular vesicle detection: A comparison of different techniques. Cytometry Part A. 2016;89(2):123–34.

57. Andreu Z, Yáñez-Mó M. Tetraspanins in extracellular vesicle formation and function. Front Immunol. 2014;5:442.

58. Muralidharan-Chari V, Clancy J, Plou C, Romao M, Chavrier P, Raposo G, D’Souza-Schorey C. ARF6-regulated shedding of tumor cell-derived plasma membrane microvesicles. Curr Biol. 2009;19(22):1875–85.

59. Hayakawa K, Chan SJ, Mandeville ET, Park JH, Bruzzese M, Montaner J, Arai K, Rosell A, Lo EH. Protective Effects of Endothelial Progenitor Cell-Derived Extracellular Mitochondria in Brain Endothelium. Stem Cells. 2018;36(9):1404–10.

60. Yamamoto H, Itoh N, Kawano S, Yatsukawa Y-i, Momose T, Makio T, Matsunaga M, Yokota M, Esaki M, Shodai T, Kohda D, Hobbs AEA, Jensen RE, Endo T. Dual role of the receptor Tom20 in specificity and efficiency of protein import into mitochondria. Proceedings of the National Academy of Sciences of the United States of America. 2011;108(1):91–6.

61. A. Naveena H, Bhatia D. Hypoxia Modulates Cellular Endocytic Pathways and Organelles with Enhanced Cell Migration and 3D Cell Invasion**. ChemBioChem.n/a(n/a):e202300506.

62. Liao L-X, Zhao M-B, Dong X, Jiang Y, Zeng K-W, Tu P-F. TDB protects vascular endothelial cells against oxygen-glucose deprivation/reperfusion-induced injury by targeting miR-34a to increase Bcl-2 expression. Scientific Reports. 2016;6(1):37959.

63. Smolina N, Bruton J, Kostareva A, Sejersen T. Assaying Mitochondrial Respiration as an Indicator of Cellular Metabolism and Fitness. Methods Mol Biol. 2017;1601:79–87.

64. Rose S, Frye RE, Slattery J, Wynne R, Tippett M, Pavliv O, Melnyk S, James SJ. Oxidative stress induces mitochondrial dysfunction in a subset of autism lymphoblastoid cell lines in a well-matched case control cohort. PLoS One. 2014;9(1):e85436.

65. Zemskova M, Kurdyukov S, James J, McClain N, Rafikov R, Rafikova O. Sex-specific stress response and HMGB1 release in pulmonary endothelial cells. PLoS One. 2020;15(4):e0231267.

66. Conchinha NV, Sokol L, Teuwen L-A, Veys K, Dumas SJ, Meta E, García-Caballero M, Geldhof V, Chen R, Treps L, Borri M, de Zeeuw P, Falkenberg KD, Dubois C, Parys M, de Rooij LPMH, Rohlenova K, Goveia J, Schoonjans L, Dewerchin M, Eelen G, Li X, Kalucka J, Carmeliet P. Protocols for endothelial cell isolation from mouse tissues: brain, choroid, lung, and muscle. STAR Protocols. 2021;2(3):100508.

