## Supplementary data for "Mitochondria-containing extracellular vesicles from mouse *vs*. human brain endothelial cells for ischemic stroke therapy"

600 Forbes Avenue, 453 Mellon Hall,

Pittsburgh, PA 15282.

X/Twitter: @manickam_lab

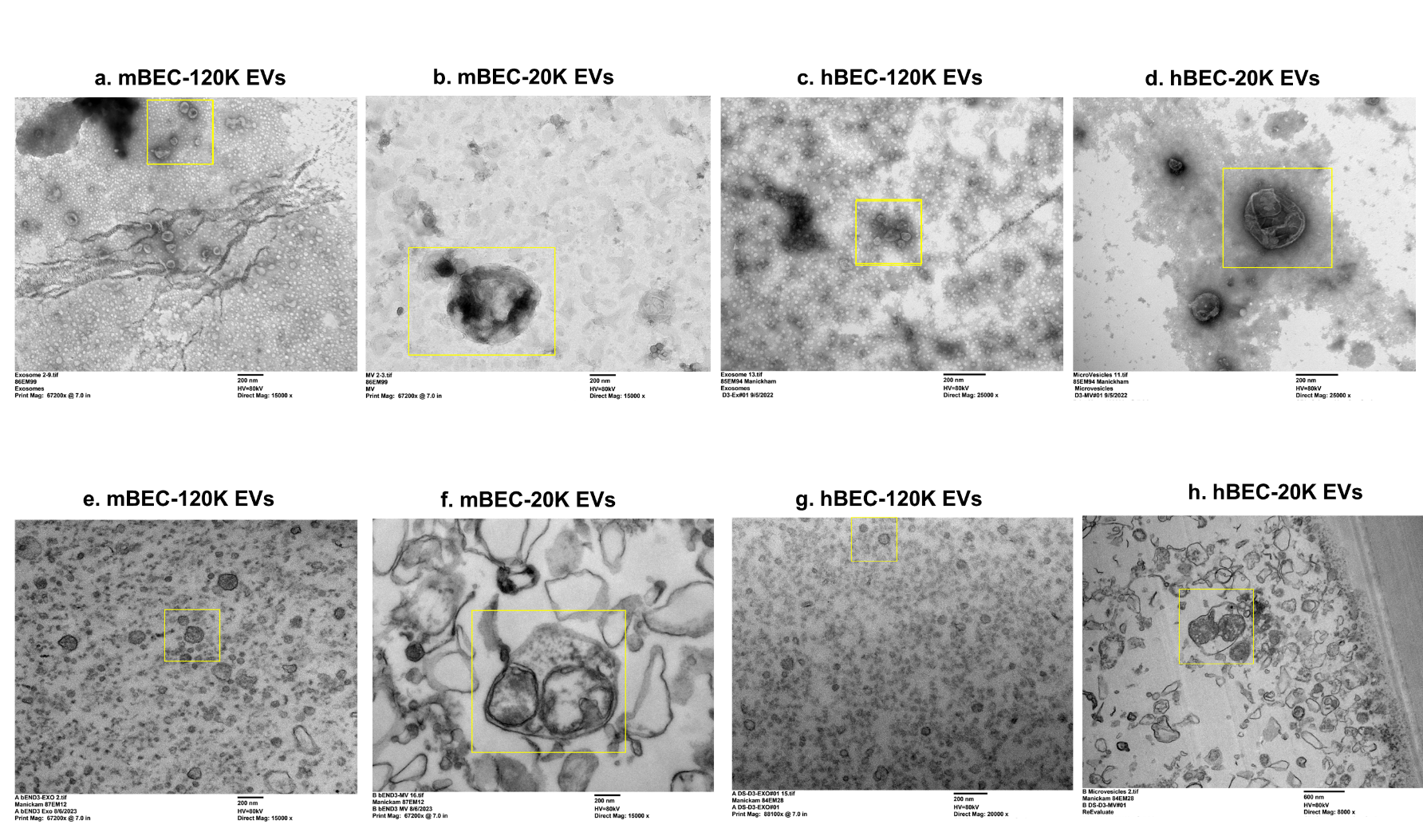

**SL Figure 1:** Uncropped TEM images of negatively-stained mBEC-120K EVs **(a)**, mBEC-20K EVs **(b)**, hBEC-120K EVs **(c)** and hBEC-20 K EVs **(d)**. Uncropped TEM images of cross-sectioned mBEC-120K EVs **(e)**, mBEC-20K EVs **(f)**, hBEC-120K EVs **(g)** and hBEC-20 K EVs **(h)**.

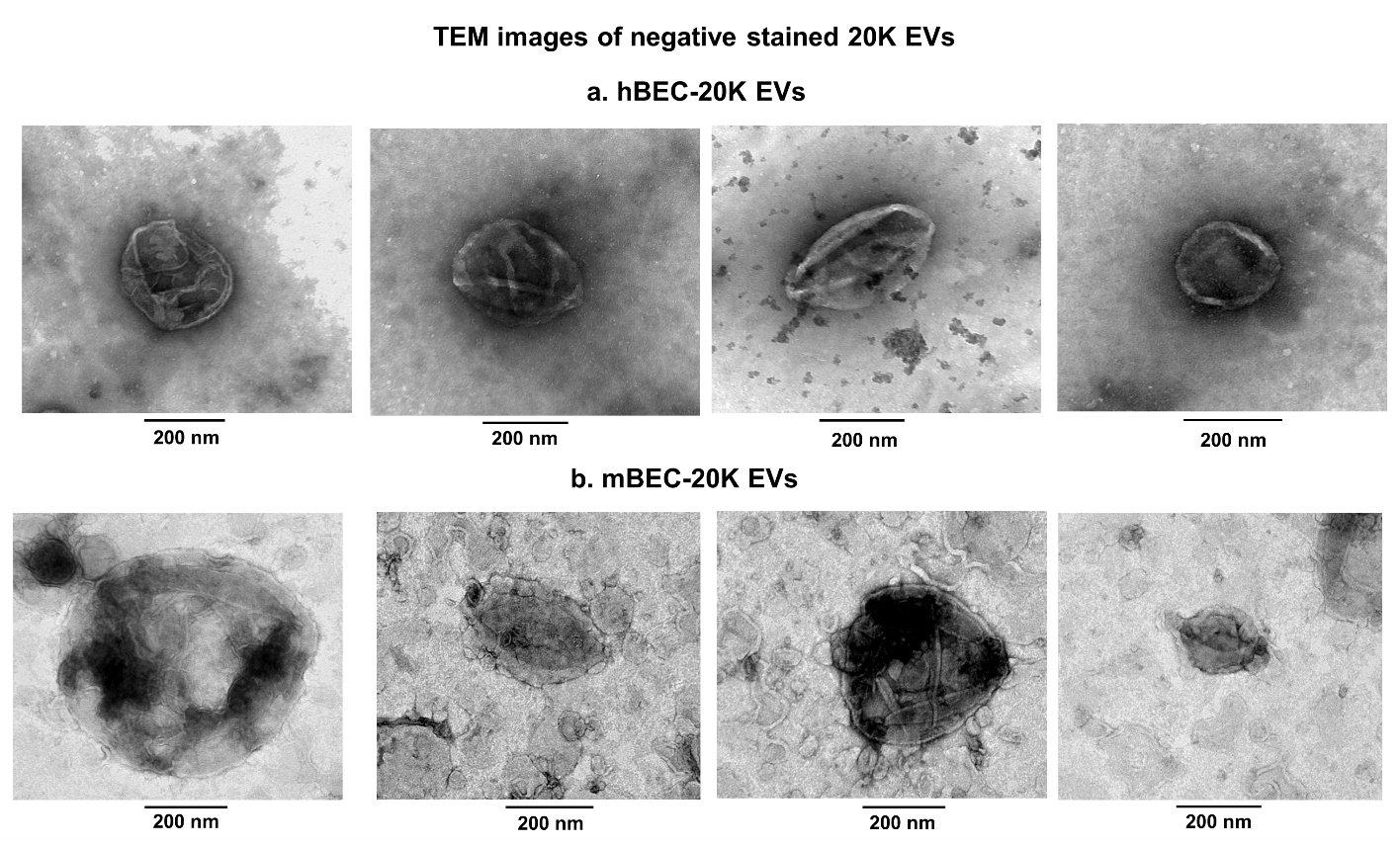

**SL Figure 2:** TEM images of negatively-stained 20K EVs derived from hBECs **(a)** and mBECs **(b).**

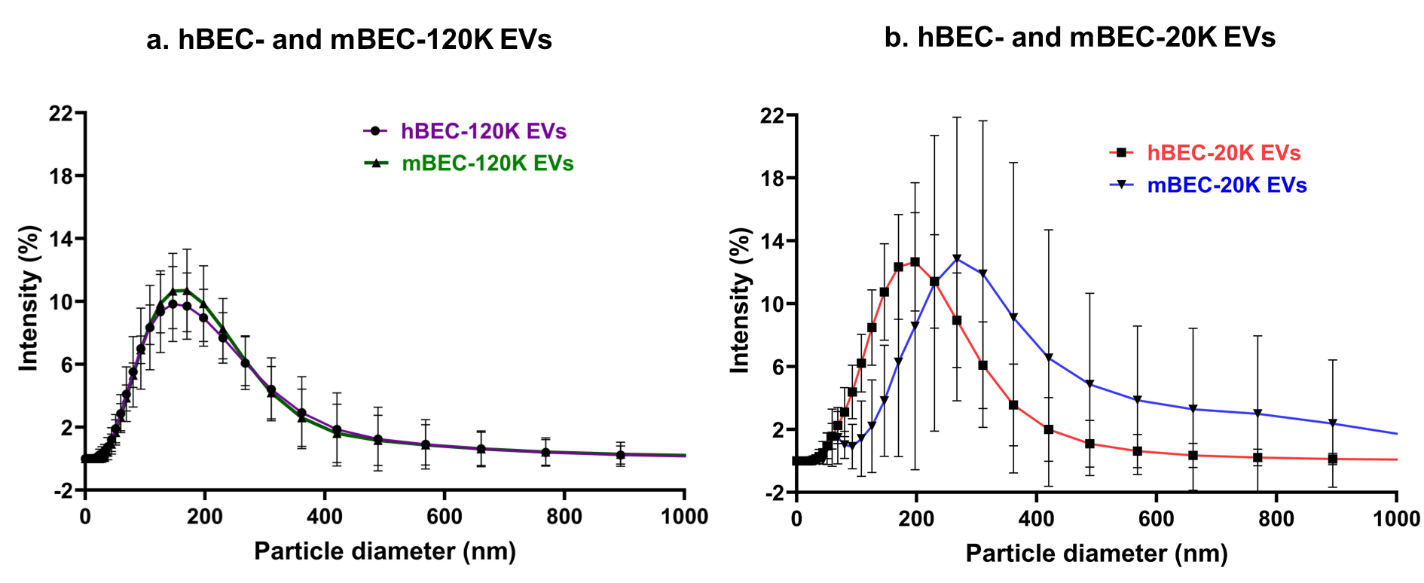

**SL Figure 3:** Intensity-weighted particle size distribution of hBEC and mBEC-derived 120K-EVs **(a)** and 20K EVs **(b)**.

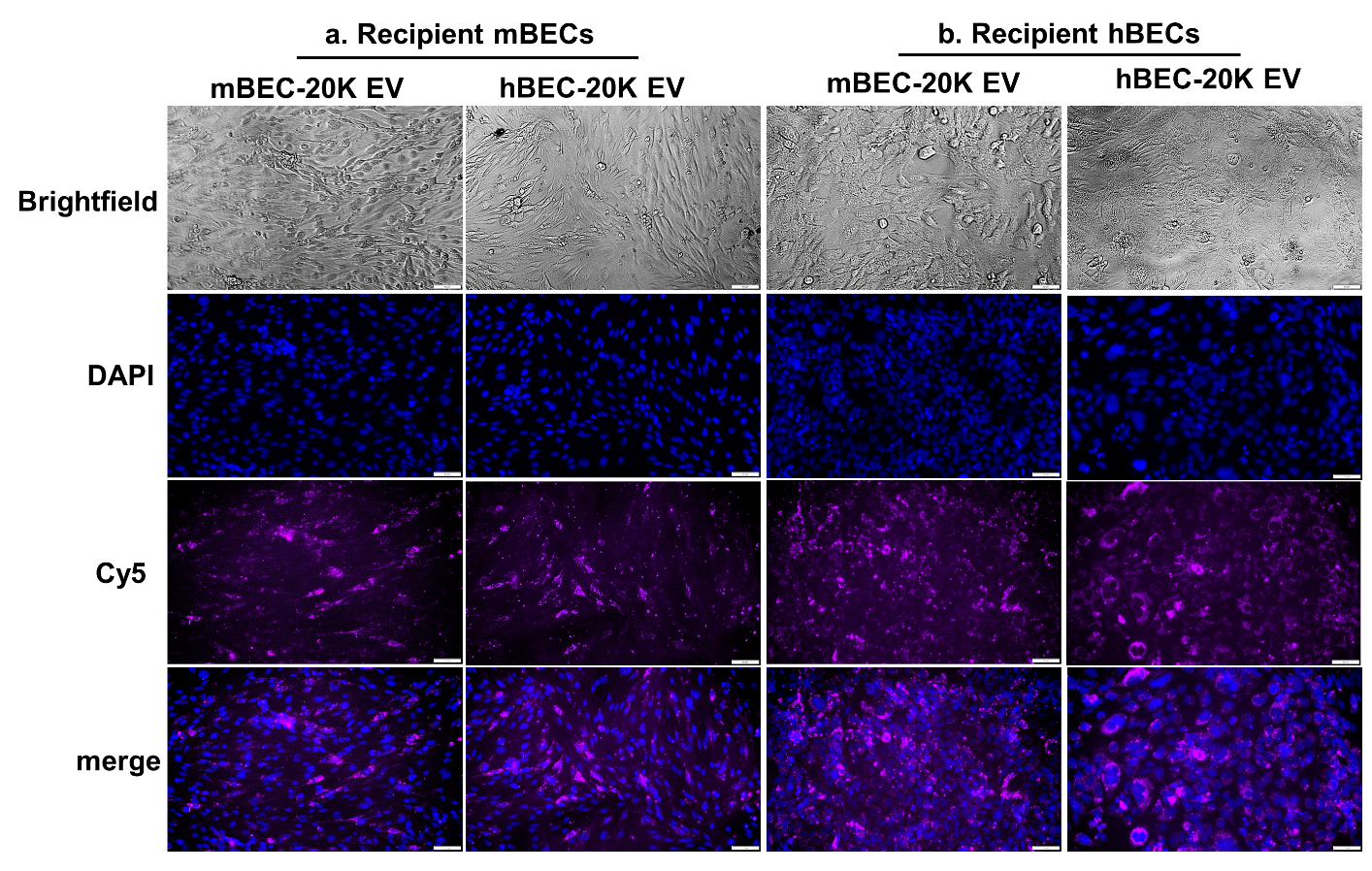

**SL Figure 4:** Uncropped raw images of recipient mBEC (bEnd.3 cell line) and hBEC (hCMEC/D3 cell line) treated with 30 μg/well with mBEC- and hBEC-derived MitoT-red-20K EVs under normoxic conditions. Representative images were acquired under brightfield, DAPI, and Cy5 channels using Olympus epifluorescent microscope . Scale bar 50 μm.

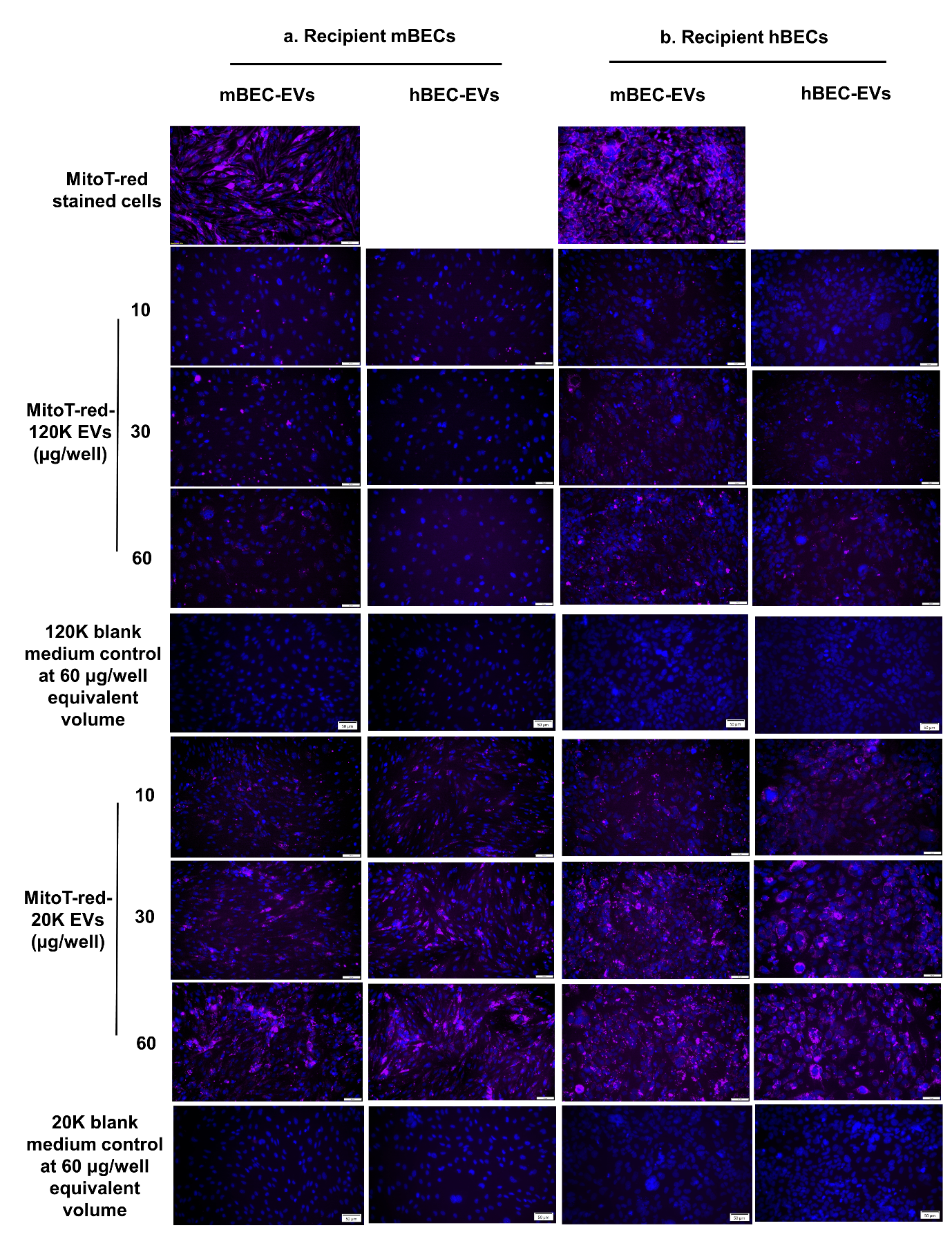

**SL Figure 5:** Uncropped raw images of recipient mBECs (**a**) and hBECs (**b**) treated with indicated concentrations of mBEC- and hBEC-derived MitoT-red-120K EVs, 20K EVs, and blank medium controls under normoxic conditions. Representative images were acquired under brightfield, DAPI, and Cy5 channels using Olympus epifluorescent microscope. Scale bar 50 μm.

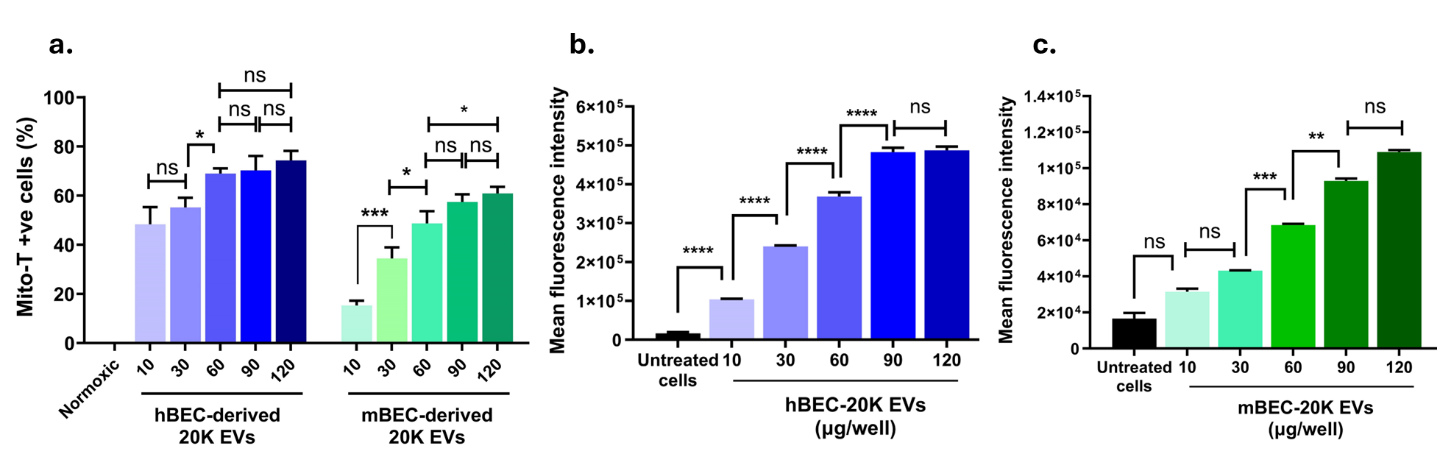

**SL Figure 6: Effect of 20K EV doses on MitoT-red-EV mitochondria transfer to recipient hBECs under normoxic conditions. (a)** % MitoT+ve recipient hBECs treated with hBEC- and mBEC-derived MitoT-red-20K-EVs at indicated doses for 48 h. **(b)** Mean fluorescence intensity (**MFI**) of hBECs treated with hBEC-derived MitoT-red-20K-EVs at indicated doses. **(c)** MFI of hBECs treated with mBEC-derived MitoT-red-20K-EVs at the indicated doses. Data represents mean±SD (n=3). * p<0.05, ** p<0.01, *** p<0.001, **** p<0.0001.

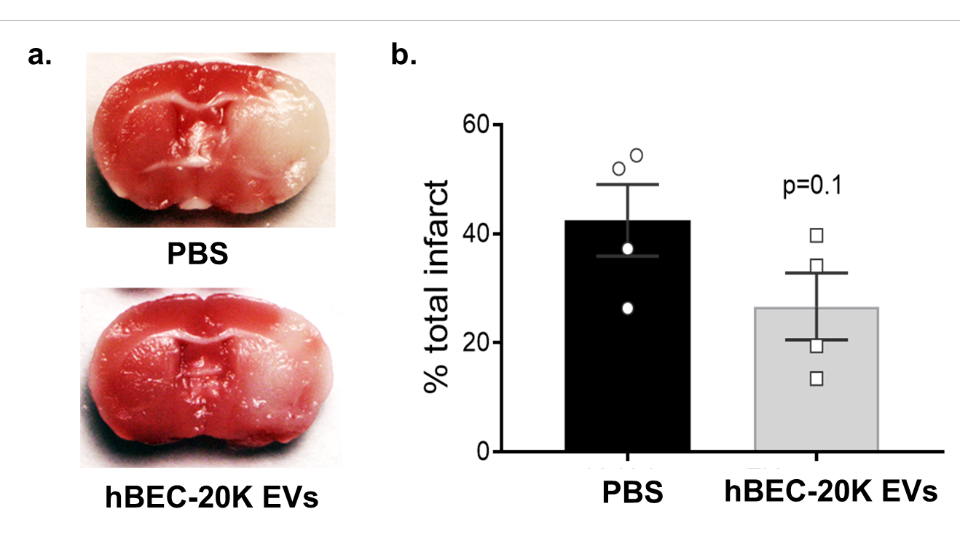

**SL Figure 7: Pilot study demonstrating potential neuroprotective effects of hBEC-20K EVs in a mouse middle cerebral artery occlusion (MCAo) model of ischemia/reperfusion injury (stroke).** **(a)** representative 2,3,5-triphenyl tetrazolium chloride (**TTC**)-stained coronal sections of the vehicle (PBS) and hBEC-20K EVs-treated stroke brains from young male mice. **(b)** Quantification of total hemispheric infarct volume at 24 h post-stroke. Data are mean ± SEM (n = 4) and were analyzed using an unpaired t-test. The figure is slightly adapted from Dave et al. *J Con Rel*. 2022 [11].
